# Network-based cytokine inference implicates Oncostatin M as a driver for inflammation phenotype of knee osteoarthritis

**DOI:** 10.1101/2023.01.24.525463

**Authors:** Hirotaka Iijima, Fan Zhang, Fabrisia Ambrosio, Yusuke Matsui

## Abstract

Inflammatory cytokines released by the synovium after trauma disturb the gene regulatory network and have been implicated in the pathophysiology of osteoarthritis. A mechanistic understanding of how aging perturbs this process can help identify novel interventions. Here, we introduced network paradigms to simulate cytokine-mediated pathological communication between the synovium and cartilage. Cartilage-specific network analysis of injured young and aged murine knees revealed aberrant matrix remodeling as a transcriptomic response unique to aged knees displaying accelerated cartilage degradation. Next, network-based cytokine inference with pharmacological manipulation uncovered IL6 family member, Oncostatin M, as a driver for the aberrant matrix remodeling. By implementing a phenotypic drug discovery approach, we identified that the activation of Oncostatin M recapitulated “inflammation” phenotype of knee osteoarthritis and highlighted high-value targets for drug development and repurposing. These findings offer translational opportunities targeting the inflammation-driven osteoarthritis phenotype.

**GRAPHICAL ABSTRACT:** 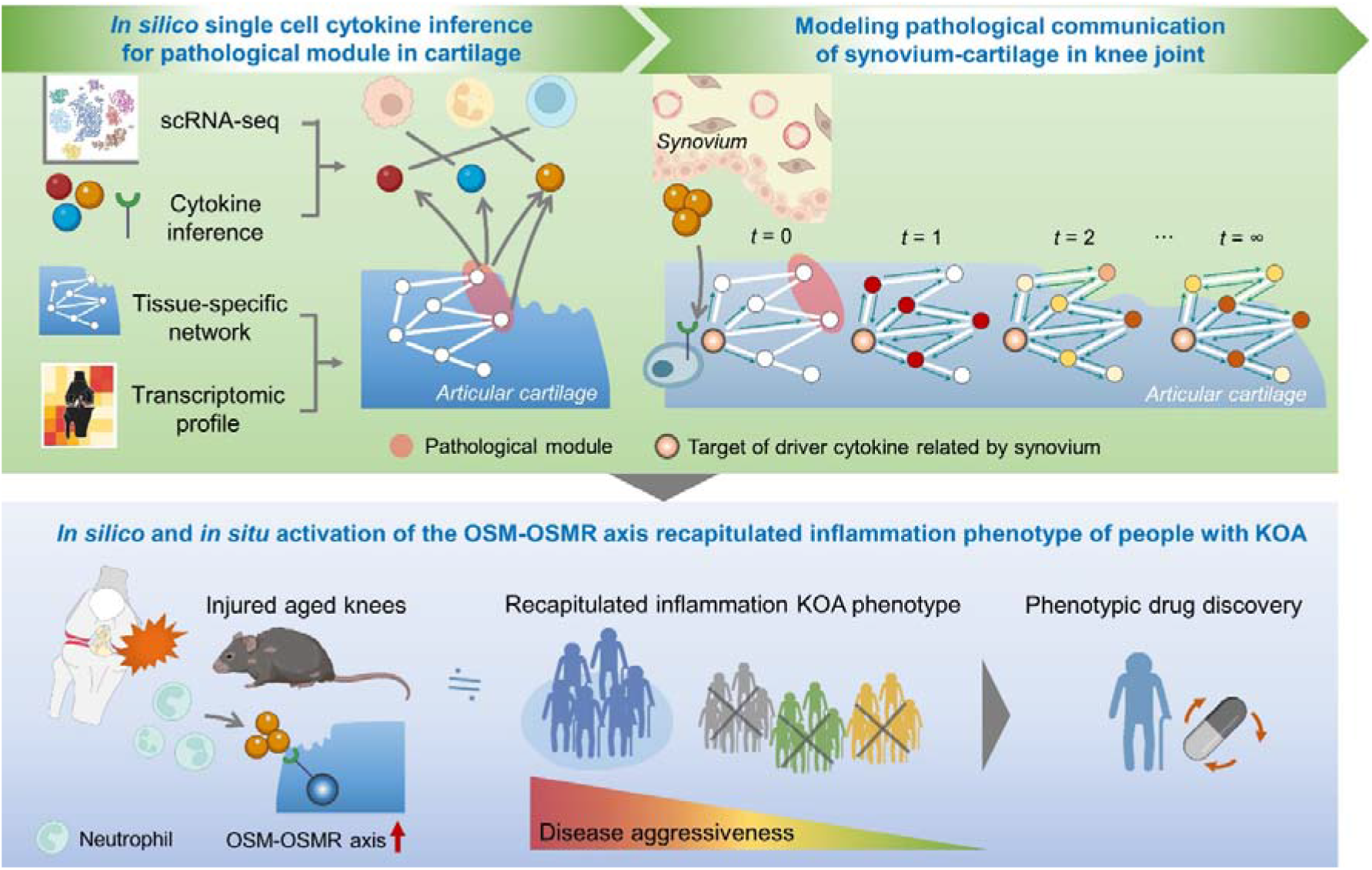

## INTRODUCTION

Adaptation, or ‘adaptive homeostasis’, is a highly conserved process, wherein cells, tissues, and whole organisms transiently activate a suite of signaling pathways in response to short-term external perturbations, thereby effecting transient changes in gene expression and stress resistance^1^. Mounting evidence suggests, however, that the capacity for adaptive homeostasis declines with aging. In 2014, Kennedy *et al*, suggested seven pillars of aging, one of which was “adaptation to stress”, whereby reduced stress capacity over time is a major risk factor for most late-onset human diseases^2^. Given the modifiable nature of stress, leveraging the relationship between stress and tissue health represents a promising area for the development of therapeutic interventions to treat aging diseases.

One of the most debilitating age-related diseases is osteoarthritis (OA) of the knee joint (KOA), which is characterized by an age- and/or mechanical stress-dependent progressive loss of cartilage integrity^3-5^. Age-associated cellular changes in articular cartilage include cell depletion due to various forms of cell death together with dysfunctional intrinsic cellular repair responses to mechanical loading^6^. This impaired intrinsic cellular repair response with aging amplifies the deleterious effects of traumatic injury^7-9^, for example, through increased levels of extracellular matrix (ECM) degrading enzymes (e.g., MMP13^10^) and decreased cartilage ECM production^9,10^. While these age-related stress responses after trauma are known accelerants of KOA, most studies investigating the transcriptomic response *in vivo* have utilized relatively young animals (∼6 months). Moreover, deleterious mechanical loading through disruption of knee structures and tissues is more common in elderly individuals than has been previously appreciated^11,12^. As a field, we therefore lack both a holistic understanding of the complex KOA disease process as well as age-dependent disease drivers that may serve as promising therapeutic targets in a geriatric population.

While KOA has historically been considered a disease of “wear and tear”, it is increasingly viewed to be a whole joint disease that results from complex tissue-tissue interactions^13^, and particularly, the synovium-cartilage crosstalk^14^. Inflammatory immune mediators have been identified in the context of inflamed synovium from rheumatoid arthritis^15,16^, but it is unclear which cytokines released from the synovium interact with cartilage in KOA. One hypothesis is that inflammatory cytokines released by the synovium disturb the complex gene regulatory network in articular cartilage and have been implicated in the pathophysiology of KOA^17^. To unravel these complex interactions, network-based cytokine inference offers a novel platform with the potential to guide further experimental work designed to uncover disease mechanisms and cytokine therapeutic targets^18^. This network medicine approach postulates a “disease module hypothesis”, in which disease-associated genes or proteins likely share the same topological neighborhood in a network^18^. Network propagation, which simultaneously considers all possible paths between given cytokine target genes and the disease-associate genes, is, therefore, a powerful tool for elucidating cytokine drivers as novel therapeutic targets. The network-based cytokine inference integrated with network propagation-based disease driver discovery is further strengthened by considering tissue specificity to understand complex molecular interactions in the manifestation of diseases^19^.

In this study, we used a series of network medicine approaches to thoroughly characterize age-related alterations in the transcriptomic responses to traumatic injury in the knee joint. We started by performing a meta-analysis of the existing literature to summarize the current knowledge of mechanisms underlying the predisposition of aged knees to accelerated KOA following trauma. Subsequent network-based cytokine inference integrated with network propagation predicted Oncostatin M and OSM receptor (OSMR) as primary cytokine drivers of aberrant ECM remodeling and stiffening with aging. These *in silico* findings were then cross-checked through systems biological analysis of archived RNA-seq data for pharmacological manipulation of OSM to murine chondrocytes *in vitro*. The culmination of these analyses suggests that aberrant ECM remodeling induced by the OSMR drives age-related accelerated KOA after trauma through pathological synovium-cartilage crosstalk. To enhance the clinical relevance of our work, these murine findings were compared with the transcriptomic signature of people with KOA and identified that activation of OSM-OSMR axis drives inflammation phenotype of KOA, the major phenotype that represents 16-30% of KOA^20^. These data suggest that targeting the OSM-OSMR axis may represent a promising new therapeutic target in the treatment of inflammation phenotype of KOA. Finally, we propose novel drug candidates that reduce OSM-OSMR signaling and reverse inflammation, offering translational opportunities for the treatment of KOA in the clinic.

## RESULTS

### Aging aggravates trauma-induced compromised cartilage integrity

While studies using cartilage explant models have suggested that aging amplifies the deleterious effects of traumatic injury^7,9^, no comprehensive study to date has summarized age-related changes in the stress response of trauma to the knee joint *in vivo*. Therefore, we first sought to address this knowledge gap by performing a systematic review of structural and molecular changes about the knee joint after traumatic injury in both young and aged mice. Six studies met the pre-specified inclusion criteria^21-26^ (**Figure 1A**). All identified studies used C57BL/6 background mice (**Table S1**). The experimental models used included destabilization of the medial meniscus (DMM), anterior cruciate ligament (ACL) transection or rupture, or tibial compressive loading (**Figure 1B**), all of which produce excessive mechanical loading to the knee joint^27^.

**Figure 1.**
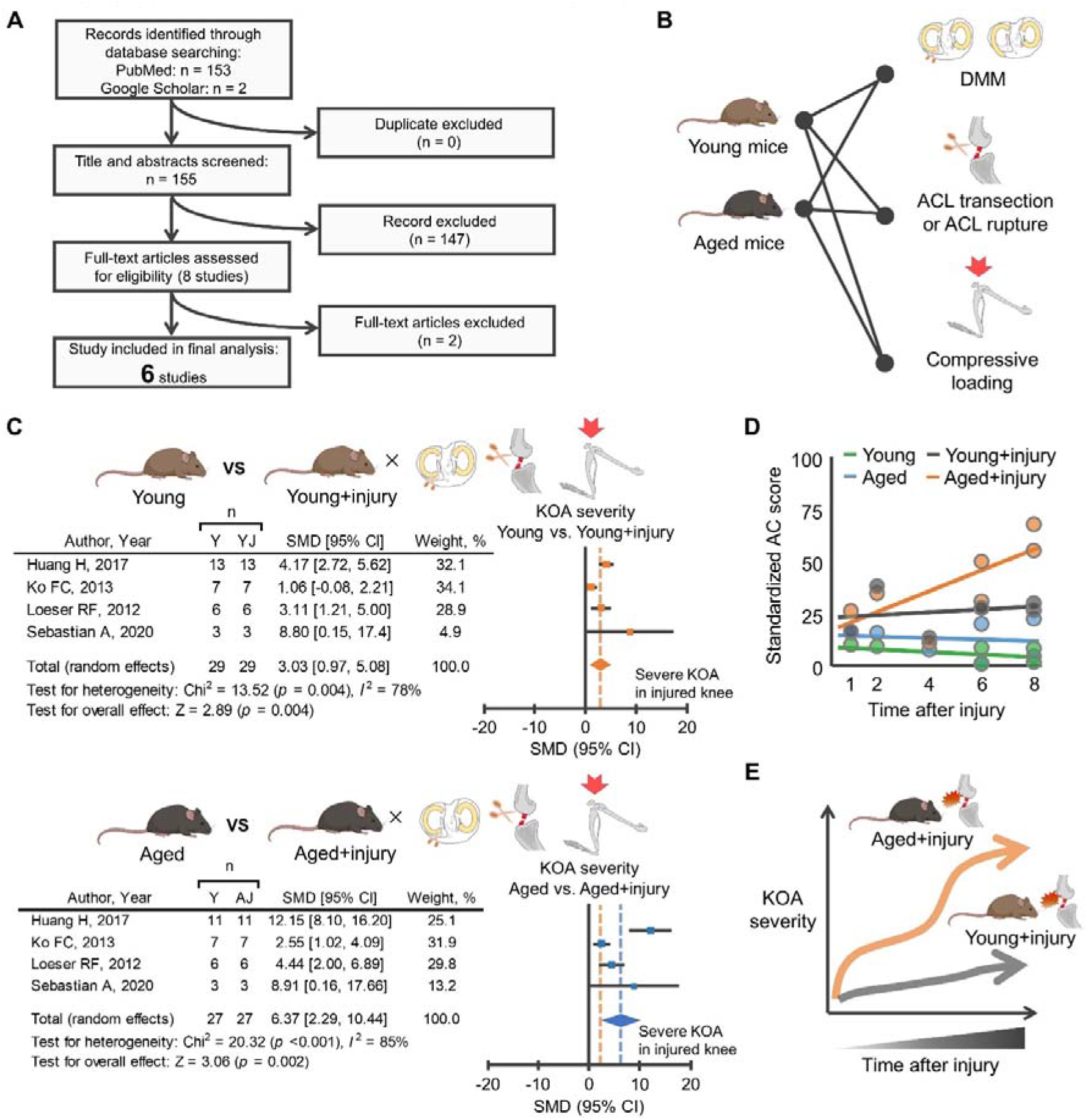
Aged mice displayed accelerated cartilage degeneration after traumatic injury. **A,** An electronic database search yielded a total of 155 studies, of which 6 studies were finally included in the meta-analysis for histological analysis. **B,** Mouse models used in the included studies were DMM, ACL transection, ACL rupture, or excessive compressive loading, all of which were used to produce mechanical overloading to knee joint. **C,** Meta-analysis of histological analysis for articular cartilage revealed that mechanical overload induces cartilage degeneration in which aged mice displayed greater cartilage degeneration compared to young mice. The forest plot displays relative weight of the individual studies, SMDs, and 95% CIs. Diamonds indicate the global estimate and its 95% CI. The orange dotted line in the lower panel indicates the average SMD of young mice. **D,** A trajectory in cartilage morphology (standardized AC score) across the four different groups (young, aged, young+injury, and aged+injury) revealed that aged, but not young, mice displayed progressive cartilage degeneration after injury. Each dot represents an individual study. **E,** Schematic illustrating the trajectory difference in KOA severity between aged and young mice after injury. Portions of the figures were created with Biorender.com. *Abbreviation: A, aged; AC, articular cartilage; AJ, aged+injury; ACL, anterior cruciate ligament; 95% CI: 95% confidence interval; DMM, destabilized medial meniscus; KOA, knee osteoarthritis; Y, young; YJ, young+injury; SMD*, *standardized mean difference*.

Subsequent meta-analysis of histological metrics confirmed that aging provokes a more severe cartilage degeneration after traumatic injury when compared to young counterparts (**Figure 1C**). For these studies, cartilage degeneration was defined as disruption of cartilage integrity assessed by semi-quantitative histological scoring. Notably, aged mice displayed progressive cartilage degeneration after traumatic injury up to eight weeks after surgery, a trend that was not observed in young mice (**Figure 1D**). The trauma-induced structural changes in the aged knee are supported by non-pooled data (i.e., data that were not included in the meta-analysis of histological metrics because of methodological heterogeneity). For example, we found that ACL transection in aged mice resulted in elevated mRNA levels of p16, a biomarker of chondrocyte aging^28^, in the knee joint when compared to young counterparts^29^.

Given that epidemiologic and biologic studies have suggested a link between cartilage degeneration and subchondral bone abnormal remodeling^30,31^, we also searched for evidence of increased subchondral bone alterations in aged knees after traumatic injury. Meta-analysis from three studies revealed that excessive mechanical loading to aged knees contributed to increased subchondral bone thickness, a surrogate marker of abnormal bone remodeling^32^, when compared to young counterparts (**Figure S1A, B**). These studies indicate that the aged joint displays accelerated degenerative changes in the cartilage-subchondral bone unit after traumatic injury relative to young counterparts (**Figure 1E**). Collectively, we interpret these data to suggest that aging exacerbates trauma-induced KOA *in vivo*.

### Cartilage-specific network analysis identified aberrant ECM remodeling as a transcriptomic signature of injured aged knees

As a next step towards a more comprehensive view of the age-related accelerated KOA after injury, we assessed the global transcriptomic changes over time following a trauma. Only one study to date has performed RNA-seq on whole-joint tissue samples from different age groups over time after trauma^26^. This study, reported by Sebastian et al^26^, demonstrated that ACL rupture results in significant alterations in genes associated with ECM remodeling and cartilage/bone metabolism in both young and aged mice relative to uninjured controls. These transcript-level findings are consistent with the disrupted cartilage integrity and increased subchondral bone thickness shown in our meta-analysis of histological metrics (**Figure 1C-E**, **Figure S1**). However, a direct comparison in the transcriptomic response to ACL rupture across the different aged mice was not performed.

To better understand age-related changes in transcriptomic responses underlying the age-related accelerated KOA, we accessed the archived RNA-seq data from young (3 months) and aged (15 months) mice from the study described above^26^. We evaluated expression data of 2,738 genes across different time points (day1, week1, week2, and week6) after ACL rupture (**Figure 2A**). We then assessed the difference in transcriptomic response (i.e., log2 fold change) across the two groups (young+ACL rupture vs. aged+ACL rupture) over time using principal component analysis (PCA). PCA of the time-course (day1 to week6) of the transcriptomic response revealed a clear segregation of age groups between weeks 1 and 2 (**Figure 2B**). Since 1-2 weeks after injury represents a phase of active tissue remodeling in which the knee joints in aged mice display disrupted cartilage-subchondral bone integrity (as shown in **Figure 1D**), we focused on the 241 overlapping differentially expressed genes (false discovery rate <0.05) that were upregulated at week1 and week2 after injury in aged, but not in young, mice (**Figure 2C**). The complete list of 241 genes is provided in **Table S2**. We labeled these 241 genes “*age-related stress response genes*”. Of note, only 14 genes were significantly upregulated at week1 and week2 after injury in young, but not aged, mice. As such, this study did not perform subsequent analyses on the young-related 14 stress response genes.

**Figure 2.**
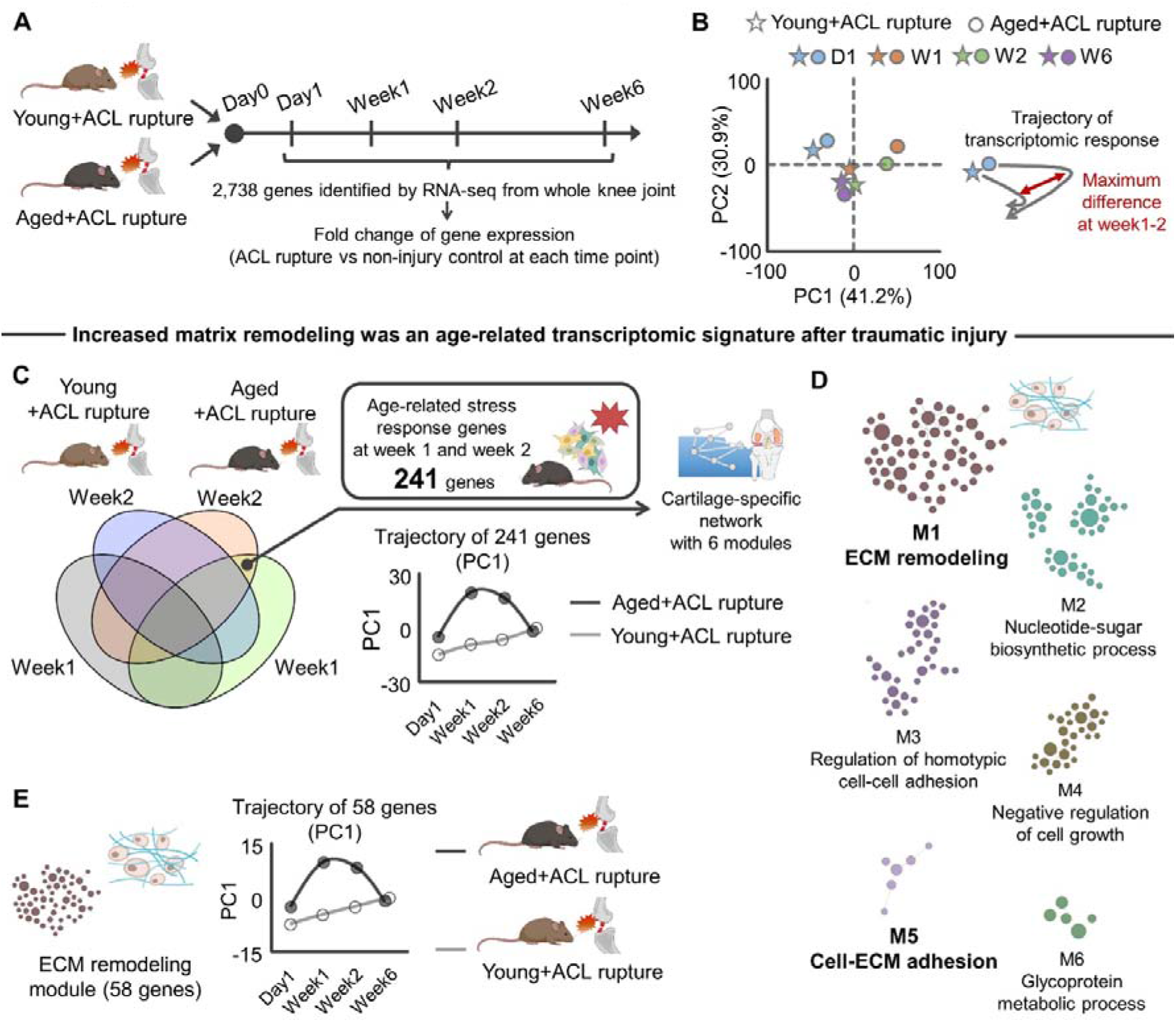
Aberrant ECM remodeling was a transcriptomic signature of aged joint after traumatic injury. **A,** Schematic showing the experimental flow of archived RNA-seq data^26^. In total, 2,738 genes were identified across the different time points (day1, week1, week2, and week6) after ACL rupture in young (3 months old) and aged (24 months old) C57/BL6 mice^26^. **B,** PCA showing the separate trajectory of time course changes (D1, day1; W1, week1; W2, week2; W6, week6) in the transcript profiles of young and aged mice with the largest different at week1 and week2. **C,** We identified 241 age-related stress response genes that were significantly upregulated in aged+ACL rupture across the different time point (week1 and week2) but not in young+ACL rupture. PCA revealed PC1 with 241 genes displayed different trajectories between young and aged mice after ACL rupture with the largest difference at week1 and week2. From 241 genes, a cartilage-specific network was constructed using HumanBase software^33^. **D,** The cartilage-specific network included six modules with the largest module annotated to “ECM remodeling”. Another small module was also annotated to “Cell-ECM adhesion”. In the cartilage-specific network, each node and edge represent gene and gene-gene interaction, respectively. **E,** PCA revealed that PC1 of ECM remodeling module genes (58 genes) displayed a different trajectory between young and aged mice after ACL rupture, with the largest difference at week1 and week2. Portions of the figures were created with Biorender.com. *Abbreviation: ACL, anterior cruciate ligament; ECM, extracellular matrix; PC, principal component; PCA, principal component analysis*.

We next sought to characterize the biological function of the age-related stress response genes identified. For this purpose, we generated a cartilage-specific functional gene network using the HumanBases software, which is a genome-scale protein function and interaction map of human tissues derived from integrating data sets from thousands of experiments^33^ (**Figure 2C**). A functional relationship implies that both genes participate in the same biological process in a specific tissue or functional context^19^. The tissue-specific functional gene network clarifies biological functions of given genes to help identify disease driver genes that may exert important functional roles according to the tissue of interest^19,34,35^.

In total, six modules were detected from the cartilage-specific network, with the biggest module (58 genes) representing “ECM remodeling”. Accordingly, “Cell-ECM adhesion” represented another module that emerged (**Figure 2D**). Of interest was the ECM remodeling module, which displayed distinct trajectories over time across the two age groups (**Figure 2E**). These alterations are in line with a previous study demonstrating that mechanical overloading to elderly osteochondral explants significantly changed the gene ontology (GO) biological process, focal adhesion^7^, a biological function directly triggered by ECM remodeling^36^. We note that the ECM remodeling module was not identified as the primary biological function when using either traditional GO enrichment analysis (Enrichr software^37^; **Figure S2**) or non-cartilage-specific (i.e., lung, bone, and heart) network analysis^33^ (**Figure S3**). This observation highlights the significance of considering the tissue-dependent network topology in interpreting biological function of genes.

Increased ECM remodeling was also identified on the cartilage-specific network using a seccond set of transcriptomic data of aged murine knee joint at 8 weeks after DMM induction^8^ (**Figure S4**), suggesting overlapping transcriptomic responses associated with ECM remodeling across different trauma models. In addition, we found that 726 genes upregulated after ACL rupture in both young and aged murine knee joints were also significantly associated with ECM remodeling (**Figure S5A-B**). For all of these, the aged knee joint displayed a greater transcriptomic response relative to young counterparts (**Figure S5C**). Among the genes of interest, we identified *Lox* as significantly upregulated by ACL rupture in both young and aged knee joints (**Figure S5D**). The protein product of *Lox*, lysyl oxidase (LOX), is one of the major enzymes involved in collagen crosslinking^38^ and has been shown to contribute to pathogenesis of post-traumatic KOA^39^. The collective findings from histological and transcriptomic analyses indicate that traumatic injury induces ECM remodeling in murine knee joints, and these transcriptomic responses are further exacerbated by aging.

Given that transcripts were collected from whole joint tissue of the knee joint, we also asked whether ECM remodeling after ACL rupture in aged mice is associated with articular chondrocyte markers. Since there are no established chondrocyte-specific markers, we defined a set of articular chondrocyte markers using archived single cell RNA-seq data from adult murine knee joints^40^. In the single cell RNA-seq data, 42 genes were predominantly expressed in articular chondrocytes (**Table S3**). Using this set as a signature, we found that the 241 age-related stress response genes were significantly enriched in chondrocytes (odds ratio: 5.96; **Figure S6A**). These data suggest that aberrant remodeling of the ECM after injury is primarily driven by aged articular chondrocytes.

### Aging amplified the trauma-induced transcriptomic responses to cartilage ECM stiffening

ECM remodeling alters biophysical properties of the ECM, often resulting in increased stiffness over time^41^. For this reason, matrix alterations have recently been proposed to be a hallmark of aging^42^. It is well established that increased ECM stiffness disrupts chondrocyte functionality via mechanotransductive signals^39,43-45^. We therefore sought to dissect biophysical from biochemical effects of the ECM. Specifically, we tested whether age-related ECM remodeling genes upregulated after ACL rupture in aged mice are associated with biophysical ECM features. For this purpose, we first defined genes associated with cartilage ECM stiffness in murine knees, as presented in published studies. For this, we again performed a systematic literature search to collect articles that examined cartilage ECM stiffness using atomic force microscopy or similar measurement systems designed to assess biophysical properties of cartilage in mice with and without gene manipulations (**Figure 3A**). In these analyses, no restriction was applied for the target gene selection. If a genetic modification significantly changed cartilage ECM stiffness, then we defined that gene as associated with cartilage stiffness.

**Figure 3.**
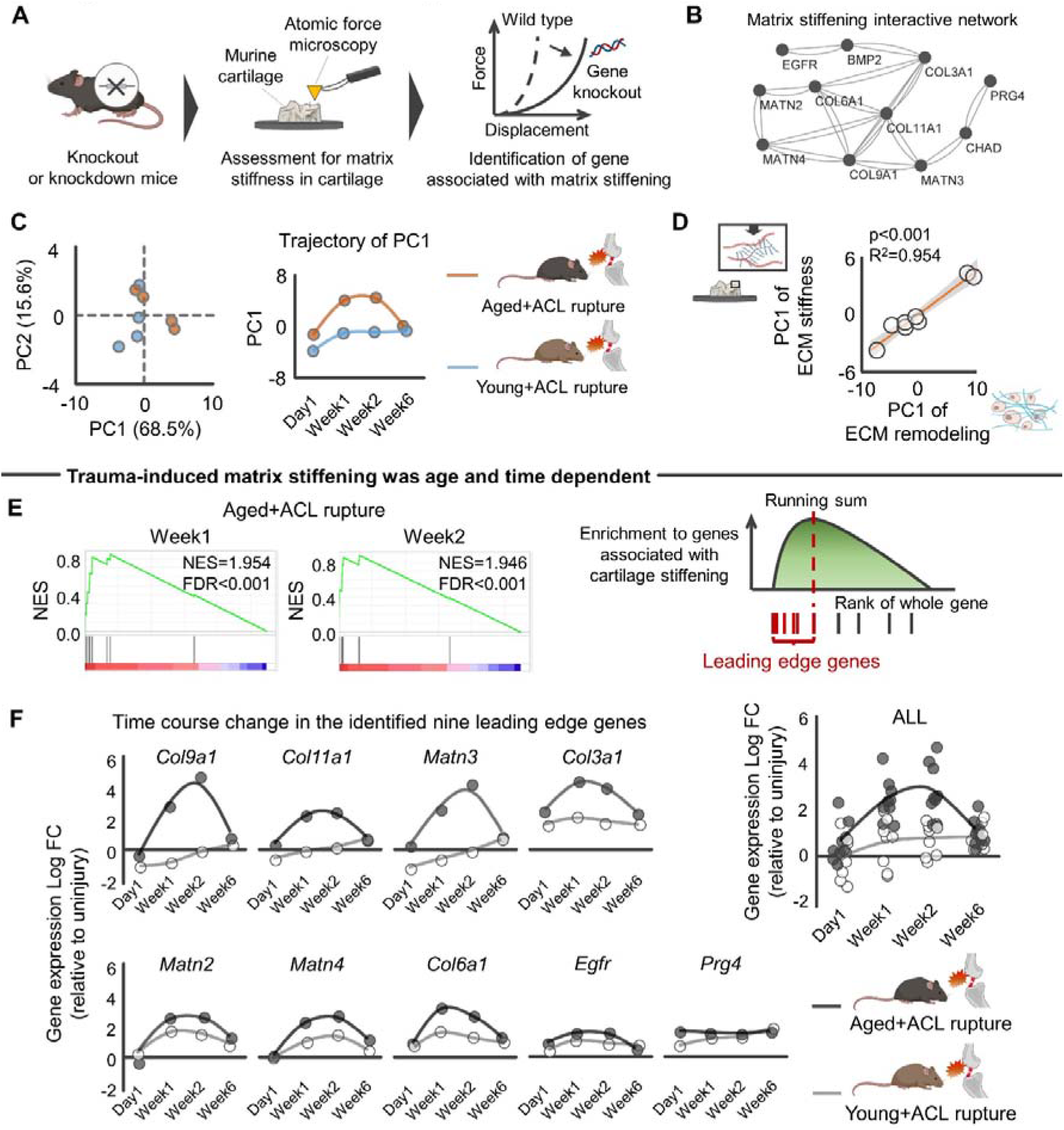
Aging-specific transcriptomic response to mechanical overload was associated with ECM stiffening in murine knee joint. **A,** Schematic showing definition of genes associated with cartilage stiffening. A systematic search identified studies assessing ECM stiffness of murine cartilage with and without gene manipulation using atomic force microscopy. **B,** ECM stiffening-related genes contributed to protein interactive network constructed by String^65^. **C,** PCA showing the separate trajectory of time course changes in transcripts response of ECM stiffening genes to ACL rupture between young and aged mice. **D,** Transcriptomic response of genes related to ECM remodeling was associated with those related to ECM stiffening. **E,** ssGSEA revealed that 241 genes upregulated in aged+ACL rupture were significantly enriched to cartilage stiffening. Nine leading edge genes were identified by ssGSEA. **F,** A time course trajectory of leading edge genes stratified by age. Statistical analyses were performed using linear regression (**D**). Portions of the figures were created with Biorender.com. *Abbreviation: ACL, anterior cruciate ligament; ECM, extracellular matrix; FC, fold change; FDR, false discovery rate; NES, normalized enrichment score; PC, principal component; PCA, principal component analysis; ssGSEA, single sample gene set enrichment analysis*.

The systematic literature search finally included 19 studies^46-64^ implementing gene knockout/knockdown (**Table S4**). Of the genes of interest, we finally identified 11 genes (*Bmp2*, *Chad*, *Col3a1*, *Col6a1*, *Col9a1*, *Col11a1*, *Egfr*, *Matn2*, *Matn3*, *Matn4*, and *Prg4*) that were associated with increased cartilage stiffness (i.e., knockout/knockdown significantly decreased cartilage stiffness). All the included genes constructed a protein-protein interactive network generated by STRINGdb database^65^ (**Figure 3B**). Interestingly, *Col11a1* emerged as a hub gene of the constructed protein-protein interactive network. *Col11a1*, genetically associated with human OA in multiple joints^66^, is a gene encoding type XI collagen, which normally interacts with type II and type IX collagen to form the meshwork of collagen fibrils in articular cartilage^67^. PCA revealed that the injured aged knee joint displayed different transcriptomic responses of 11 ECM stiffness-related genes when compared to young counterparts (**Figure 3C**). As expected, the transcriptomic response of ECM stiffness-related genes was significantly associated with those of age-related aberrant ECM remodeling (**Figure 3D**). We interpret these results to suggest that the aberrant ECM remodeling unique to aged knees is, at least partly, accompanied by ECM stiffening.

To further support the identified relationship between an age-related altered transcriptomic response to traumatic injury and cartilage stiffening, we performed single sample gene set enrichment analysis (ssGSEA)^68^ with transcriptomic response of aged knees at week1 and week2 after ACL rupture^26^, in which the 11 genes associated with increased ECM stiffness were included as a gene set. An extension of gene set enrichment analysis (GSEA)^68^, ssGSEA, calculates separate enrichment scores for each paring of a sample and gene set. The goal of ssGSEA was to prioritize genes significantly correlated with a given phenotype (i.e., ECM stiffening). Using this analysis, we found that ACL rupture in the aged knee increased expression of genes associated with increased cartilage stiffness, in which nine genes (*Col3a1*, *Col6a1*, *Col9a1*, *Col11a1*, *Matn2*, *Mant3*, *Matn4*, *Prg4*, and *Egfr*) were identified as leading-edge genes that were maximally upregulated at week1 and week2 after ACL rupture in aged, but not young, mice (**Figure 3E-F**). These results suggest that aging amplifies trauma-induced ECM stiffening.

### Network-based cytokine inference predicted the OSM-OSMR axis as a driver of age-related aberrant ECM remodeling

We next tested for candidate drivers of the observed aberrant ECM remodeling with aging. Accumulating evidence has shown that ECM remodeling is induced by inflammatory cytokines^69^. Traumatic knee injuries trigger an immediate increase of inflammatory cytokines in the synovial fluid^70^. Since inflammatory synovial cytokines disturb joint homeostasis and are implicated in the pathophysiology of KOA^17^, we explored cytokines released exclusively by the synovium as potential upstream regulators for the age-related aberrant ECM remodeling in the aged knee after injury (**Figure 4A**). Using the 58 genes related to ECM remodeling identified above as inputs (**Figure 2D-E**), CytoSig analysis^71^ identified a total of 43 cytokines, in which 12 cytokines were predicted to be elevated in the aged knee joint after ACL rupture relative to young counterparts (**Figure 4A**). Of these 12 cytokines, Oncostatin M (OSM) and IL6 were predominantly expressed in synovial cells but not in chondrocytes (<1%), according to single cell RNA-seq data of articular cartilage and synovium isolated from elderly individuals with KOA (GSE152805)^14^ (**Figure 4B**).

**Figure 4.**
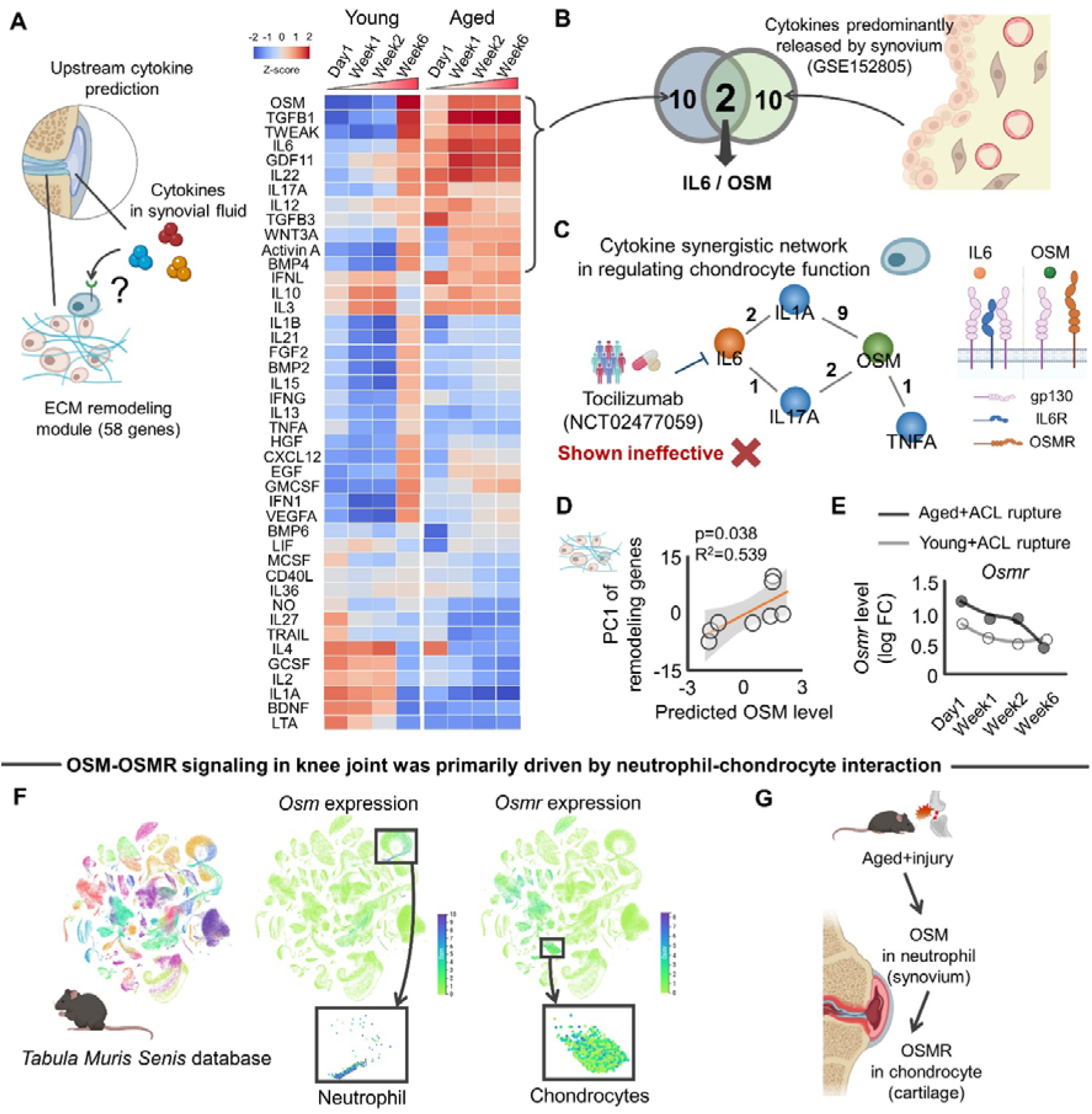
Activated OSM-OSMR was predicted driver for age-related aberrant ECM remodeling. **A,** Predicted upstream cytokines driving the age-related aberrant ECM remodeling in the cartilage-specific network analyzed by CytoSig software^71^. Heatmap color indicates z-score for each gene. Of 43 cytokines, 12 were predominantly upregulated in aged knee joints. **B,** Integrated analysis of upstream cytokine inference and archived bulk RNA-seq of synovium (GSE152805)^14^ identified IL6 and OSM as possible cytokine drivers predominantly released by the synovium. **C,** Constructed synergistic cytokine network that regulates ECM remodeling in chondrocytes (left panel). Numbers in the network represent the number of studies supporting the synergistic effect (p-value <0.05). Results of clinical trial (NCT02477059) targeting IL6 is embedded in the network for illustration purpose. Ligand-receptor complex of IL6 as well as OSMR are also provided for illustration purpose. **D,** Predicted OSM level was significantly associated with injury-induced transcriptomic response of ECM remodeling genes. **E,** Trajectory of *Osmr* gene expression after ACL rupture. **F,** *Tabula Muris Senis* database revealed that while *Osm* expression was predominantly driven by neutrophils, *Osmr* expression was predominantly driven by chondrocytes. **G,** Schematic illustrating possible neutrophil-chondrocyte interaction that activates the OSM-OSMR axis in aged murine knee joints after mechanical overload. Statistical analyses were performed using linear regression (**D**). Portions of the figures were created with Biorender.com. *Abbreviation: ACL, anterior cruciate ligament; ECM, extracellular matrix; PC, principal component*.

IL6 and OSM are members of the IL6 family of cytokines and are produced by immune cells in response to tissue injury^72^. IL6 signaling is actively involved in regulating cartilage integrity and pain, implicating IL6 as a potential therapeutic target for murine post-traumatic knees^73^. However, a recently conducted clinical trial of IL6 receptor antagonist, Tocilizumab, showed no significant pain relief in older subjects with hand OA when compared to placebo^74^. OSM is a promising alternative to IL6 agonists because OSM contributes to the network of cytokines, mediators, enzymes, and structural proteins that control ECM homeostasis, which are unique to OSM among the IL6 family^75^. OSM primes and amplifies inflammatory responses to IL1, TNF-α, or IL17A in joint^75^, all of which lead to compromised cartilage integrity. Indeed, when we originally defined the synergistic network of cytokines surrounding to OSM and IL6 in regulating cartilage ECM remodeling (e.g., TIMPs and MMPs) through systematic literature search^76-87^, OSM served as a local hub of the cytokine synergistic network (**Figure 4C**). Full information for the synergistic effects of cytokines is available in **Table S5**. Further supporting the regulatory role of OSM, the predicted OSM level was positively associated with age-related remodeling genes (**Figure 4D**).

While OSM and IL6 have redundant biological functions due to the common signal transducing receptor, gp130, OSM also produces non-redundant signals through its unique receptor, OSMR^72^. Since signals downstream of OSMR mediate OSM-induced ECM remodeling^88^, we revisited the individual transcriptomic responses of the archived RNA-seq data^26^. We found that OSMR was significantly increased in aged mice up to 2 weeks after ACL rupture (false discovery rate <0.001; **Figure 4E**), displaying a trajectory similar to the predicted OSM level. These findings are in line with previous studies demonstrating that patients with KOA displayed increased OSM concentration in synovial fluid and that osteoarthritic cartilage displayed increased OSMR levels^89,90^. Notably, an KOA-related increase in OSM within the synovial fluid inhibits ECM synthesis by chondrocytes, an effect that was counteracted by inhibiting OSM^89^. Although OSM also binds to Leukemia inhibitory factor receptor (LIFR) according to the comprehensive ligand-receptor database, CellChat^91^, LIFR was not detected in the RNA-seq data^26^. Together, these findings indicate that OSM may be a driver of pathogenic ECM remodeling in aged cartilage via the OSM-OSMR axis.

In light of the catabolic effect of OSM-OSMR shown in previous studies, we next sought to determine the cellular origin of these age-dependent responses to mechanical overloading. For this purpose, we accessed the *Tabula Muris Senis* database^92^ to assess which cells predominantly express OSM and OSMR in murine tissue. Results revealed that, whereas OSM is predominantly expressed in immune cells, and especially neutrophils, OSMR is predominantly expressed in chondrocytes (**Figure 4F**). These findings from murine studies are in line with the human study which demonstrated that, while OSM is highly expressed in HLA-DRA^+^ cells (including immune regulatory cells, inflammatory macrophages, dendritic cells, activated proinflammatory fibroblasts, and B cells), OSMR is highly expressed in chondrocytes in the synovium and/or cartilage samples in people with KOA^14^. Neutrophils are the first immune cells to be recruited in the inflamed joint, where they secrete proinflammatory mediators^93^. Since articular cartilage is comprised exclusively of chondrocytes, these data indicate neutrophil-to-chondrocyte interactions emanating from the synovium and cartilage, respectively. This potential neutrophil-to-chondrocyte interaction is in line with a previous study demonstrating that patients with KOA displayed increased neutrophil-related inflammation in the synovial fluid^94,95^. Exaggerated neutrophil recruitment contributes to the development of tissue damage and fibrosis^69^, which may be partly triggered by IL-17^96^, one of the cytokines significantly predicted by our cytokine inference analysis (**Figure 4A**). Indeed, a recent study showed that IL-17 levels in inguinal lymph nodes of aged mice after traumatic knee injury were significantly higher than those in young mice, while IL-17 neutralization reduced traumatic injury-induced cartilage degeneration in aged mice^29^. Taken together, these data led us to the novel hypothesis that a trauma-induced increase in OSM from the synovium binds to OSMR in chondrocytes, initiating downstream signal activation, age-related aberrant ECM remodeling, and ultimately, chondrocyte dysfunction (**Figure 4G**).

### *In silico* and *in situ* activation of the OSM-OSMR axis induced aberrant ECM remodeling in aged chondrocytes

To test the above hypothesis suggested, we implemented network propagation applied to the cartilage-specific gene network. This approach is based on the premise that network propagation represents tissue interaction of OSM (released by synovium) and OSMR (chondrocyte in articular cartilage) (**Figure 5A**). Network propagation explores the network vicinity of seeded genes to study their functions based on the premise that nodes with similar functions tend to lie close to each other in the networks^97^. To simulate, we first constructed a knowledge-driven global cartilage-specific network using HumanBases software^33^. With this global network in hand, we posited that *in silico* activation of *Osmr* in chondrocytes would activate the age-related stress response genes we found above to be overexpressed with trauma in aged joints. For this purpose, a random-walk-restart (RWR) algorithm^98^ was used to verify which nodes are most frequently visited on a random path in a given network. As expected, RWR applied to cartilage-specific network revealed that genes pseudo-activated (i.e., an affinity score >0) by *Osmr* were significantly associated with age-related stress response genes (**Figure 5B**). This finding suggests that age-related stress response genes are in the vicinity of *Osmr* and, therefore, age-related aberrant ECM remodeling, a primary functional module of age-related stress response genes, are functionally connected with *Osmr* in regulating cartilage homeostasis.

**Figure 5.**
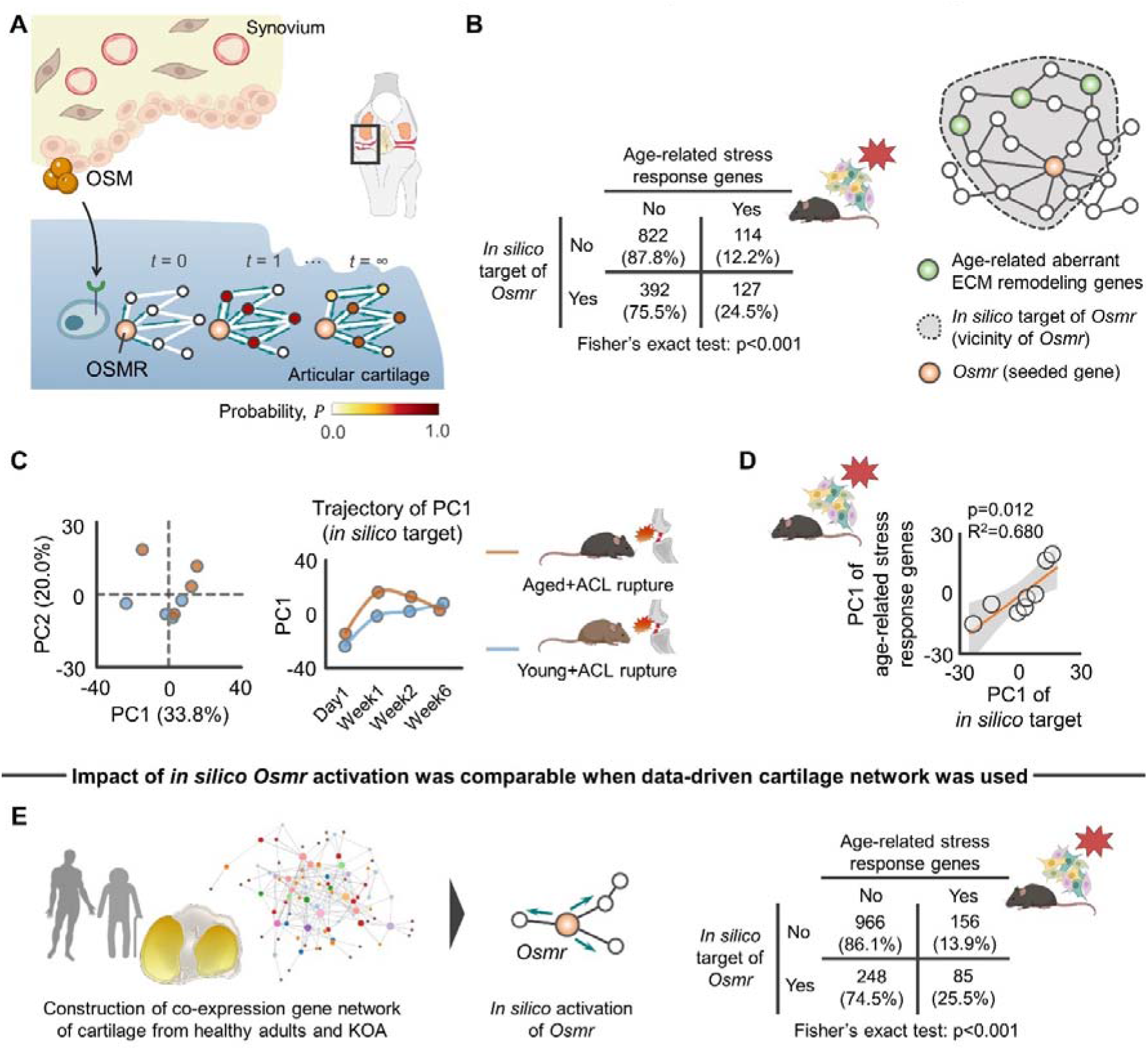
*In silico* network approach predicting the impact of pseudo activated OSM-OSMR axis for cartilage-specific network. **A,** Schematic showing tissue-specific network propagation that simulates tissue-tissue interaction of synovium (OSM) and cartilage (OSMR) in knee joints. **B,** *In silico* target genes of *Osmr* displayed significant enrichment to age-related stress response genes. Values in the 2×2 contingency table show number of genes (percentage). **C,** PCA showing the separate trajectory of time course changes in transcripts response of *in silico* target genes of *Osmr* to ACL rupture between young and aged mice. **D,** Transcriptomic response of *in silico* target genes of *Osmr* was associated with age-related stress response genes. **E,** Sensitivity analysis of *in silico* activation of *Osmr* on the different co-expression networks constructed from cartilage samples of people with healthy adults and KOA. WGCNA^101^ was used to construct the network according to bulk RNA-seq data (GSE114007)^99^.Values in the 2×2 contingency table show number of genes (percentage). Statistical analyses were performed using Fisher’s exact test (**B** and **E**) or linear regression (**D**). Portions of the figures were created with Biorender.com. *Abbreviation: ACL, anterior cruciate ligament; ECM, extracellular matrix; KOA, knee osteoarthritis; PC, principal component; PCA, principal component analysis; WGCNA, weighted gene correlation network analysis*.

To compare the transcriptomic response of *in silico* target genes of *Osmr* with age-related stress response genes against traumatic injury, we revisited archived RNA-seq data^26^ and performed PCA with the transcriptomic response of *in silico* target genes included as input. PCA revealed that the injured aged knee joint displayed a distinct transcriptomic response to *in silico* perturbation of *Osmr* target genes when compared to young counterparts (**Figure 5C**). Intriguingly, the transcriptomic response of *in silico* target genes of *Osmr* was significantly associated with age-related stress response genes (**Figure 5D**), supporting the hypothesis that activated OSM-OSMR axis recapitulates an age-related transcriptomic response to injury and induces aberrant ECM remodeling in aged cartilage.

To address the possibility that the effects of *in silico* activation on age-related stress response genes is a function of type of network constructed (i.e., knowledge-driven vs. data-driven), we also built the data-driven gene co-expression network based on the transcriptomic response of human cartilage (GSE114007)^99^. The data-driven *de novo* networks indicate putative biomolecular interactions within a specific biological condition and help to elucidate disease networks and predict therapeutics in a more holistic way^100^. Weighted gene correlation network analysis (WGCNA) is a data-driven approach to generate gene-gene co-expression networks from all pairwise correlations^101^. Consistent with findings from the original network constructed by HumanBases (**Figure 5A-B**), pseudo-activated genes by *in silico Osmr* activation were significantly associated with age-related stress response genes (**Figure 5E**).

In addition to the *in silico* analysis, we also sought to show direct evidence of OSM on age-related aberrant ECM remodeling *in situ*. We accessed archived microarray data of primary chondrocytes that were isolated from murine knee joints with or without OSM supplementation^102^ (**Figure 6A**). The original study by Liu et al was designed to compare transcriptomic response of articular chondrocytes to treatments by IL6 family cytokines (OSM, IL6, IL11, or leukemia inhibitory factor; 100ng/mL in each cytokine)^102^. The results showed that OSM supplementation significantly changed the expression level of 2,373 genes, among which OSMR was significantly upregulated after treatment (**Figure 6B**). Of note, among the 2,373 genes identified, nine genes were highly upregulated (mean expression >10, log fold change >2.5). The nine upregulated genes were significantly associated with inflammation-related biological function such as “Regulation of inflammatory response” and “Neutrophil chemotaxis” (**Figure 6B**), which is in line with our findings that OSM was predominantly released by neutrophils (**Figure 4F**).

**Figure 6.**
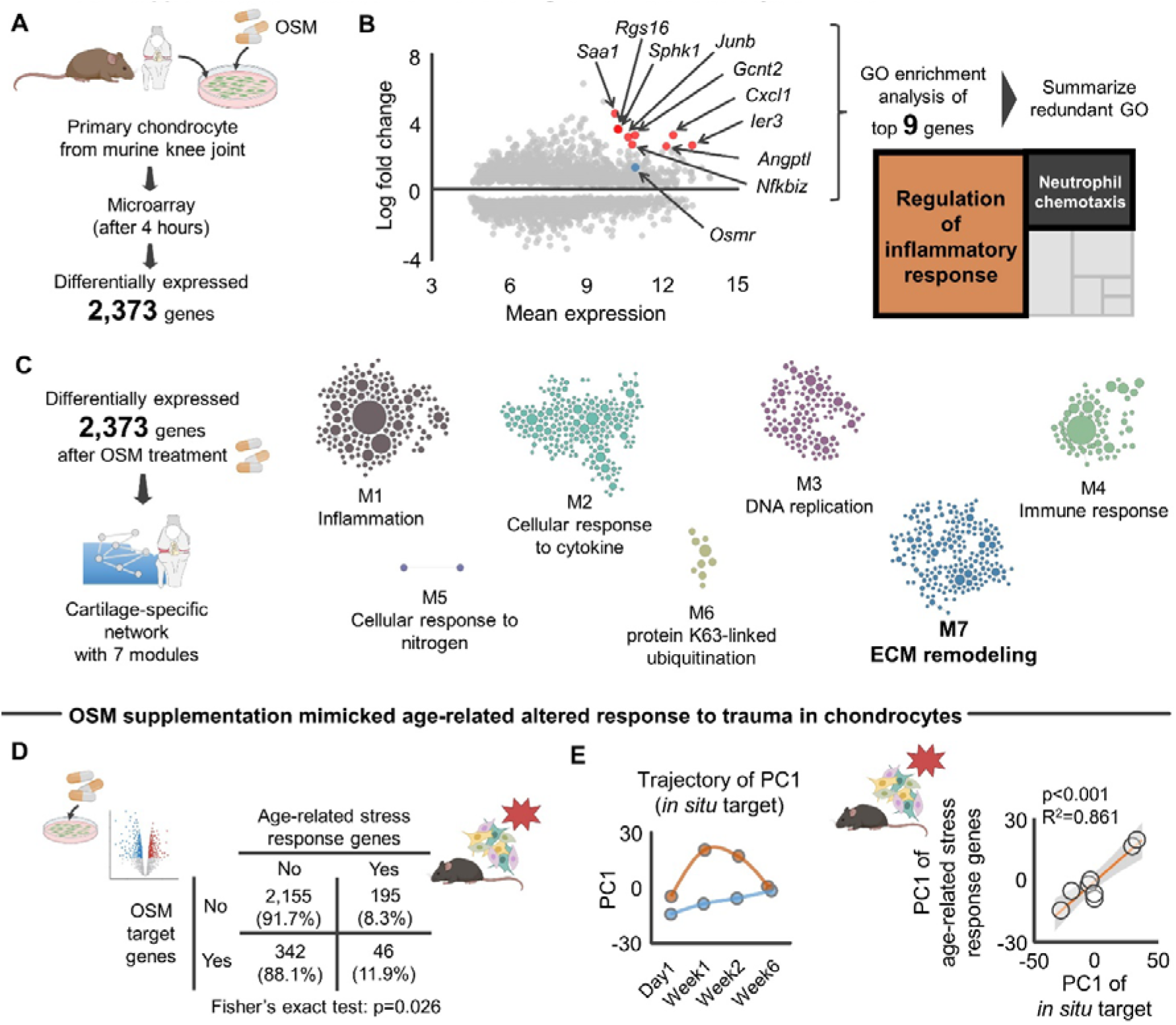
OSM influenced age-related aberrant ECM remodeling genes in murine chondrocytes. **A,** Schematic showing the experimental flow of microarray data +/- OSM supplementation to primary murine chondrocytes^102^. The microarray revealed 2,373 differentially expressed gene at 4 hours after OSM supplementation to chondrocytes. **B,** GO enrichment analysis with subsequent summarizing redundant GO of top nine genes significantly upregulated (mean expression >10, log fold change >2.5) identified inflammation-related GO biological functions, such as “Regulation of inflammatory response” and “Neutrophil chemotaxis”. In the MA plot, OSMR is highlighted with blue color for a descriptive purpose. **C,** Cartilage-specific network constructed based on 2,373 differentially expressed genes. The second largest module (module7: M7) was annotated as “ECM remodeling”. **D,** Association between differentially expressed 2,373 genes after OSM supplementation and age-related stress response genes. Values in the 2×2 contingency table indicate the number of genes (percentage). **E,** PCA showing the separate trajectory of time course changes in transcripts response of *in situ* target genes of OSM to ACL rupture between young and aged mice. Transcriptomic responses of *in situ* target genes of OSM were associated with those age-related stress response genes. Statistical analyses were performed using Fisher’s exact test (**D**) or linear regression (**E**). Portions of the figures were created with Biorender.com. *Abbreviation: ACL, anterior cruciate ligament; ECM, extracellular matrix; GO, gene ontology; PC, principal component; PCA*.

We then determined the biological function of 2,373 differentially expressed genes using a cartilage-specific network and identified an ECM remodeling module (**Figure 6C**). Most notably, the differentially expressed genes after OSM supplementation were significantly associated with age-related stress response genes (**Figure 6D**). Further, similar to our *in silico* prediction (**Figure 5C-D**), the aged knee joint displayed different transcriptomic responses of *Osmr* target genes *in situ* (**Figure 6E**), which were also significantly associated with age-related stress response genes (**Figure 6E**). Since the original study by Liu et al demonstrated the divergent transcriptomic response among the IL6 family cytokines^102^, we additionally accessed archived microarray data of primary chondrocytes that were isolated from murine knee joints with or without IL6 supplementation^102^ and asked whether the observed age-related stress response after OSM supplementation is driven by a unique biological function of OSM that is independent from IL6, another cytokine predicted to be a possible driver for the aberrant ECM remodeling (**Figure 4B**). We found that the genes significantly regulated by OSM, but not IL6, were significantly associated with age-related stress response genes (**Figure S7**). Interestingly, this significant relationship was not confirmed in genes that were significantly regulated by IL6 (**Figure S7**), indicating that OSM-induced age-related stress response after trauma in chondrocytes is predominantly driven by genes uniquely responsive to OSM.

### *In silico* and *in situ* activation of the OSM-OSMR axis recapitulated inflammation phenotype of people with KOA

Finally, we sought to correlate the findings from the murine studies with pathophysiology of KOA. Since KOA is a complex heterogeneous disease with multiple etiologies, recent studies have been dividing the KOA into different phenotypes^103^. Among the suggested phenotypes (“GAG metabolic disorder,” “collagen metabolic disorder,” “activated sensory neurons,” and “inflammation”), critical interest has been directed to “inflammation” phenotype because of therapeutic potential by anti-inflammatory drug^104^. Using single cell RNA-seq approach of human osteoarthritic cartilage, a recent study characterized the inflammation phenotype as an elevated immune response, upregulated CD34, increased crosstalk of synovium-cartilage-subchondral bone, and severe disease severity^105^ (**Figure 7A**). In addition to these findings from the original study, we found that the inflammation phenotype displays significantly higher enrichment to OSM signaling pathway (**Figure 7A**). These clinical and biological features are shared in injured age murine knees, as evidenced by elevated inflammation, structural damage of cartilage-subchondral bone, and increased *Cd34* expression compared to young counterparts (**Figure 7B**). The similarities suggest that aged murine knees after traumatic injury recapitulates the inflammation phenotypes of people with KOA.

**Figure 7.**
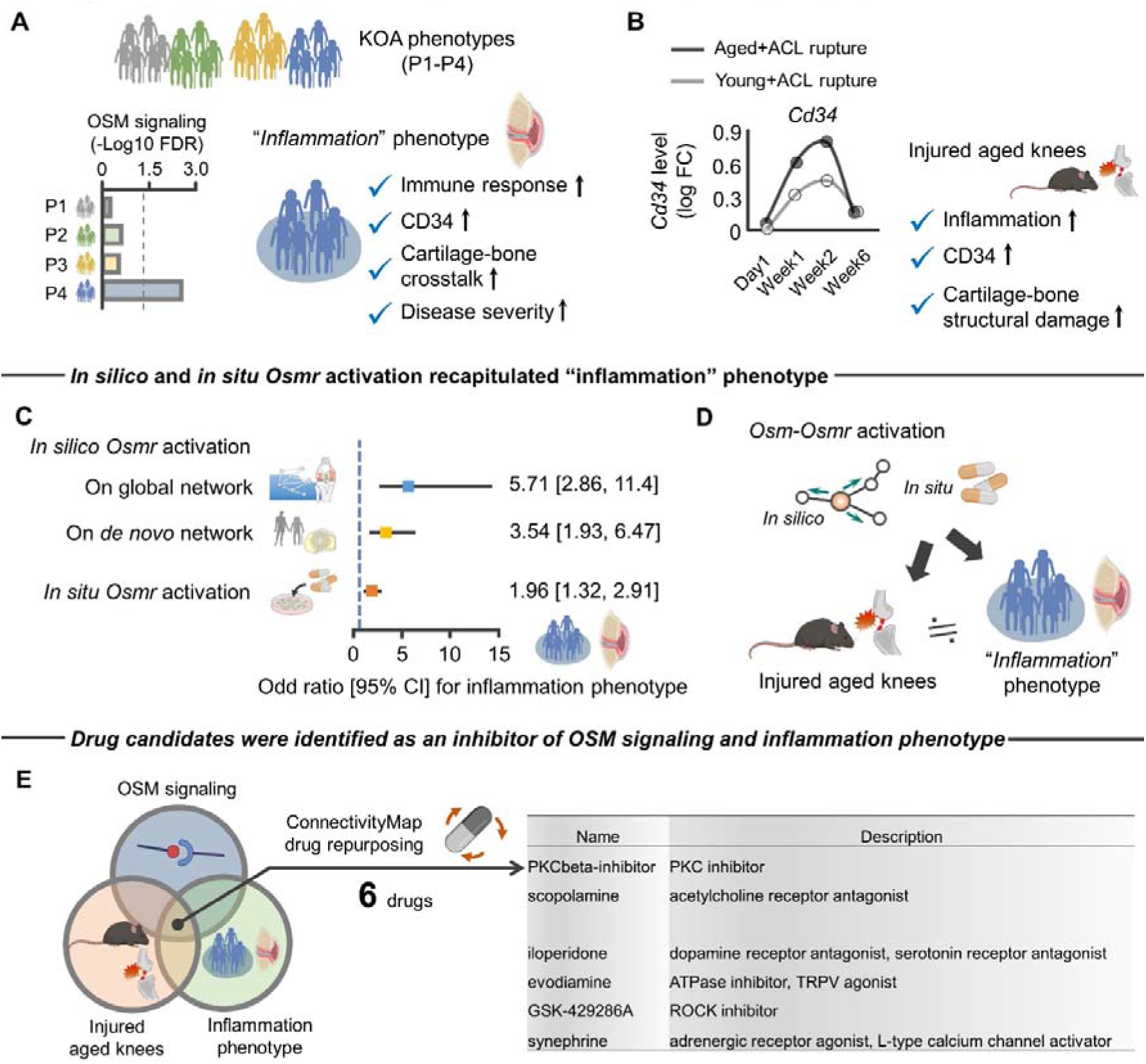
Injured aged murine knees and activation of *Osmr* recapitulated “inflammation” KOA phenotype. **A,** Schematic showing the molecular and clinical features of an “inflammation” phenotype, a one of the four phenotypes (P1-P4) of people with KOA that are defined by previous study^105^. Only the inflammation phenotype displayed significant enrichment to OSM signaling pathway. P1, GAG metabolic disorder; P2, collagen metabolic disorder; P3, activated sensory neurons; P4, inflammation. **B,** Aged mice after ACL rupture displayed increased *Cd34* gene expression compared to young counterparts. Injured aged murine knees have a similar molecular (inflammation and CD34) and structural (cartilage-subchondral bone damage) profiles seen in the inflammation phenotype of people with KOA. **C,** *In silico* and *in situ* target genes of *Osmr* displayed significant enrichment to genes associated with the inflammation phenotype (i.e., odd ratio in each model is over 1). Results were similar across the two different *in silico* networks model (cartilage-specific network constructed by Humanbases^33^ and *de novo* co-expression network constructed by WGCNA according to archived RNA-seq data (GSE114007)^99^). The blue dotted line represents an odd ratio with 1. **D,** Schematic summarizing the findings that the injured aged murine knees and activation of *Osmr* recapitulated “inflammation” KOA phenotype. **E**, Six compounds with the strongest evidence for inducing gene expression signatures that counter elevated Oncostatin M signaling, upregulated age-related stress response genes, and inflammation phenotype. All of these compounds were provided by ConnectivityMap^106^. Statistical analyses were performed using binary logistic regression analysis (**C**). *Abbreviation: ACL, anterior cruciate ligament; KOA, knee osteoarthritis; WGCNA, weighted gene correlation network analysis*.

We further simulated whether *Osmr* is a possible driver of pathophysiology of the inflammation phenotype. Using the identified set of genes (1,721 genes) that characterized the inflammation phenotype as a reference^105^, we again repeated *in silico* and *in situ* activation of *Osmr*. Consistent with the close relationship between injured aged murine knees and inflammation phenotype, *Osmr* activation significantly changed expression levels of genes associated with inflammation phenotype (**Figure 7C**). Taken together, these findings suggest that injured aged murine knees recapitulate pathophysiology of inflammation phenotype of KOA in humans, in which elevated OSM-OSMR axis may be the one possible cytokine driver (**Figure 7D**).

### Identification of potential drugs to attenuate OSM signaling and inflammation phenotype

Having molecular similarities across OSM signaling, injured aged knees, and inflammation phenotype in KOA, we sought to identify compounds capable of reversing these changes. Using in vitro drug screen data from ConnectivityMap^106^, we identified six compounds that induce strong opposing gene expression signature across the targets (i.e., OSM signaling, injured aged knees, and inflammation phenotype), attenuating the disease burden (**Figure 7E**). Some of these compounds have known biological relevance to pathophysiology of KOA, including TRPV agonist and ROCK inhibitor^39,107^ (**Figure 7E**). Intriguingly, ROCK is a known regulator of ECM remodeling^39^, consistent with our findings of age-related aberrant ECM remodeling and stiffening as a downstream of OSM signaling (**Figure 4B** and **Figure 5E**).

Further notable examples of compounds with the potential to reverse pathological molecular changes included dopamine receptor antagonist that has therapeutic potential and attenuate IL-17-mediated neutrophilic airway inflammation^108^. Other compounds, including PKCβ inhibitor and adrenergic receptor agonist, also have anti-inflammatory effects^109,110^, serving as a possible new strategy for treating inflammation phenotype of KOA. These compounds may have a candidate role in preventing the onset and/or progression of inflammation phenotype of KOA.

## DISCUSSION

An age-related decline in intrinsic chondrocyte repair responses to traumatic injury has been suggested to accelerate the progression of KOA^7-9^. With the aging of the global population and the expanding prevalence of KOA, there is a growing need to understand how aging and trauma-induced stress interact to accelerate KOA. To address this critical issue, this study introduced a network paradigm of cytokine inference integrated with network propagation in a cartilage-specific network. Using this approach, we uncovered a previously unappreciated cytokine driver, OSM-OSMR axis, for age-related aberrant ECM remodeling, possibly via inter-cellular communication of synovial neutrophils and articular chondrocytes. The aberrant ECM remodeling was accompanied by transcriptomic responses that induce ECM stiffening. These age-related altered transcriptomic responses are in line with tissue level changes, as evidenced by our meta-analysis demonstrating that aged knee joints display accelerated compromised cartilage integrity following injury when compared to young counterparts. Intriguingly, while OSM is a IL6 family cytokine, pharmacological manipulations suggested that the age-related aberrant ECM remodeling is attributed, at least partly, to downstream signals of OSM, but not IL6. The molecular and tissue level changes in injured aged knees recapitulate the pathophysiology of inflammation phenotype in people with KOA, in which the OSM-OSMR axis is a possible cytokine driver and potential novel target for the development of new therapies. The subsequent drug repurpose was built upon this theme and identified six molecules with potential clinical application to attenuate the OSM-OSMR axis as well as inflammation phenotype in people with KOA.

A critical knowledge gap in the investigation of age-related accelerated KOA has been the lack of characterization of an age-specific signature of transcriptomic in response to a traumatic injury. With the identified stress response genes unique to aged knee joints in hand, this study implemented cartilage-specific network analysis. Results revealed that aberrant remodeling and stiffening of the ECM after trauma is an age-specific feature. ECM remodeling is a widely-recognized cause of ECM stiffening^111^. In turn, ECM stiffening is a known driver of chondrocyte dysfunction and disrupted cartilage integrity via mechanotransductive pathways, both in the post-traumatic KOA in young mice^39^ as well as naturally occurring KOA in aged mice^112^. Studies have shown that traumatic injury in young murine knee joints display altered transcripts associated with ECM remodeling and induced ECM stiffening via LOX-mediated increased collagen crosslinking^26,39^. Findings of the current study suggest that aging amplifies the trauma-induced ECM remodeling and stiffening, driving an accelerated disease process.

In these analyses, the utility of implementing the cartilage-specific network analysis is supported by our finding that neither traditional GO enrichment analysis nor non-cartilage-specific network identified ECM remodeling as a primary biological function altered in aged knee joints. It should be emphasized that, owing to the low chondrocyte density (∼2% of the total volume of cartilage^113^) and the lack of blood supply in articular cartilage, chondrocyte behavior largely depends on input signals from surrounding ECM^114^. This tissue feature is unique to articular cartilage and emphasizes the need for implementation of tissue-specific network in inferring the cellular response to external environmental perturbations. The consideration of tissue specificity is much needed in the context of aging research, particularly given accumulating evidence showing that aging induces transcriptomic changes in a tissue-specific manner^115,116^.

The development and progression of KOA are now believed to involve inflammation even in the early stage of the disease^117^. Synovial cells coordinate the production of molecules that initiate and maintain synovial inflammation and contribute to cartilage damage in the setting of OA^118^. Among the proinflammatory cytokines involved in KOA, IL-1β and TNFα have been considered to be major regulators of cartilage degeneration^119^. Indeed, aged mouse knee joints displayed prolonged suppression of proteoglycan synthesis after IL-1β exposure when compared to young mice^120^. However, the results of KOA clinical trials conducted to date, which have mostly studied the effect of anti-IL1β and anti-TNF therapies, have yielded disappointing results^121^. Beyond IL-1β and TNFα, other cytokines are being investigated as targets of treatment for KOA, including IL6. IL6 targets JAK/STAT signaling, which has been implicated in the pathogenesis of inflammatory and autoimmune diseases including rheumatoid arthritis^122^. However, as was observed for IL-1β and TNFα, a recent clinical trial (NCT02477059) targeting anti-IL6 also showed no significant pain relief effect in patients with hand OA^74^. Using network-based cytokine inference integrated with network propagation that simulates synovium-cartilage crosstalk, the current study identified the OSM-OSMR axis as a predicted driver for the aberrant ECM remodeling that we found to be unique to the aged knee joint after injury. Intriguingly, pharmacological manipulation revealed that OSM-OSMR axis induces differential transcriptomic responses in chondrocytes as compared to IL6. Moreover, the genes uniquely regulated by OSM-OSMR were significantly associated with age-related aberrant ECM remodeling after traumatic injury. Secreted OSM binds its unique receptor, OSMR, which, in turn, activates downstream JAK/STAT signaling pathway to promote ECM remodeling^88,123^ and senescence of cartilage-derived stem/progenitor cells^124^. While, to our knowledge, no clinical study in KOA has been designed to target OSM, a new generation of anti-OSM antibody, GSK2330811, has been recently developed and tested in a clinical trial for the treatment of systemic sclerosis^125^.

Our network-based cytokine inference also identified IL17A as a possible driver cytokine for age-related aberrant ECM remodeling and stiffening. As indicated by the synergistic cytokine network we defined, IL17 acts an amplification factor together with OSM for other inflammatory cytokines^75,116^. IL17A activates neutrophils through production of IL8, a key chemokine for neutrophils^116^. Given that OSM originates from neutrophils, as indicated by *Tabula Muris Senis*, IL17A may be upstream of OSM release by neutrophils. While this IL17A-OSM (neutrophil) axis has been traditionally implicated in the pathogenesis of rheumatoid arthritis^116,126^, evidence has suggested it may also play a role in the onset of KOA^94,95,127^. For example, IL17A induces chondrocyte senescence and trauma-induced KOA in aged murine knees^29^. Another study showed that neutrophils may constitute a population of as much as 25% of the cells in the synovial fluid in patients with KOA^128,129^.Neutrophil elastase in synovial fluid induces chondrocyte apoptosis and activates the caspase signaling in OA^130^. Further, IL17A significantly amplifies OSM-induced production of ECM degradation enzymes in human cartilage explant and chondrocyte^131^, suggesting that therapeutics targeting IL17A together with OSM may be a promising approach to prevent or delay trauma-induced KOA in aged knees. Pre-clinical and subsequent clinical studies targeting the synergistic effects of these cytokines represent an interesting future direction.

The findings of the current study have important clinical implications for the development of novel therapeutics. Correlating the downstream effects of OSM-OSMR axis with transcriptomic signature of phenotypes in people with KOA, we discovered that activation of the OSM-OSMR axis recapitulated inflammation phenotype of KOA and identified high-value targets for drug development and repurposing. These findings offer translational opportunities targeting the inflammation-driven KOA phenotype and bring us one step closer towards establishing phenotypic drug discoveries for people with KOA.

Although this study provides a new perspective to the pathogenesis of age-related KOA, it has limitations. First, our meta-analysis for age-related accelerated KOA was based on a small number of studies, which may contribute to bias depending on the methodology used in the original studies. Additionally, given the lack of female mice in current studies, we were not able to clarify mechanisms underlying age-related KOA in female mice, and this remains an important area for future work. Finally, findings of this study were predominantly obtained from the C57BL6 mice strain. Therefore, we cannot exclude the possibility that OSM-OSMR axis as a driver of ECM remodeling may be strain specific.

While this study focused on KOA as a model, a major conceptional innovation of our work is the network-based cytokine inference integrated with network propagation on cartilage-specific network to simulate signal transduction in cartilage as a downstream of synovium-derived cytokines. It is widely recognized that age-related, low-grade inflammation –inflammaging – contributes to the pathogenesis of age-related diseases via chronic activation of the innate immune system involving inflammatory cytokines^132^. As such, we anticipate that the introduced network approaches use here may have broader applications in the field of aging research, even beyond KOA.

## Acknowledgments

This study was supported in part by (1) a JSPS KAKENHI (Grant-in-Aid for Early-Career Scientists; Grant Number: 18045240) from the Japan Society for the Promotion of Science (https://www.jsps.go.jp/) and (2) Tokai Pathways to Global Excellence Project (T-GEx) (https://www.t-gex.nagoya-u.ac.jp/en/). The funders had no role in study design, data collection and analysis, decision to publish, or preparation of the manuscript.

## Author contributions

All authors made substantial contributions in the following areas: (1) conception and design of the study, acquisition of data, analysis and interpretation of data, drafting of the article; (2) final approval of the article version to be submitted; and (3) agreement to be personally accountable for the author’s own contributions and to ensure that questions related to the accuracy are appropriately investigated, resolved, and the resolution documented in the literature.

The specific contributions of the authors are as follows:

H.I., F.Z., Y.M. provided the concept, idea and experimental design for the studies. H.I., F.A., Y.M. wrote the manuscript. H.I., F.Z., F.A., Y.M. provided data collection, analyses, interpretation and review of the manuscript. H.I. obtained funding for the studies.

## Inclusion and diversity

We support inclusive, diverse, and equitable conduct of research.

## STAR METHODS

### KEY RESOURCE TABLE

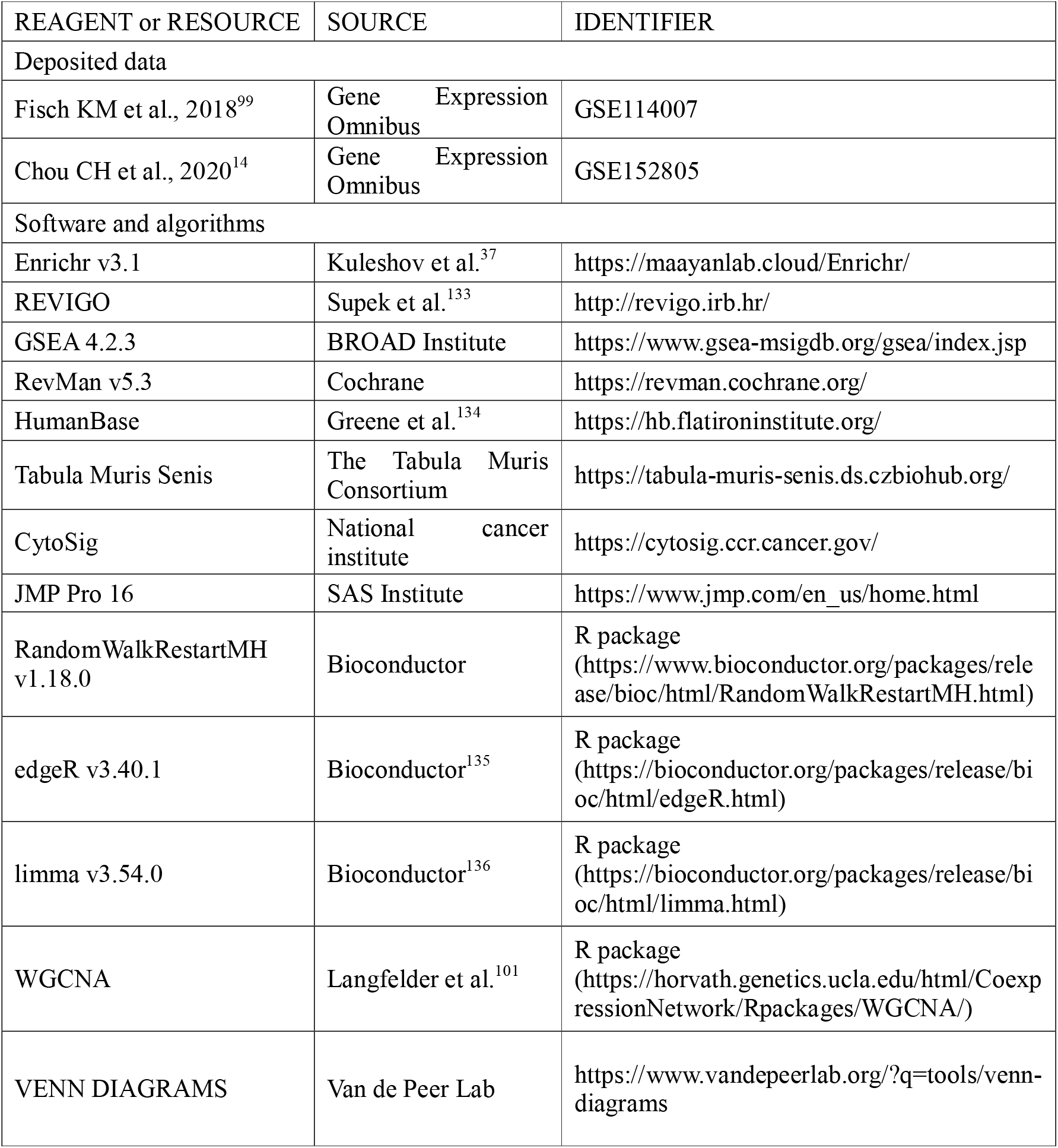

### RESOURCE AVAILABILITY

#### Lead contact

Further information and requests for resources should be directed to and will be fulfilled by the lead contact, Hirotaka Iijima.

#### Materials availability

This study did not generate new unique reagents.

#### Data and code availability

The raw data that support the experimental findings are included as Source Data. Any additional information required to reanalyze the data reported in this work is available from the lead author upon request.

### EXPERIMENTAL MODEL AND SUBJECT DETAILS

#### Animals

This study used archived data of animal experiments.

### METHOD DETAILS

#### Systematic literature search for age-related alterations to mechanical overloading

The systematic literature search was conducted according to the Preferred Reporting Items for Systematic reviews and Meta-Analyses (PRISMA) statement^137^, PRISMA protocols (PRISMA-P)^138^, Meta-analysis of Observational Studies in Epidemiology (MOOSE) checklist^139^, Cochrane Handbook for Systematic Reviews of Interventions^140^, and the practical guide for meta-analysis from animal studies^141^.

Manuscript eligibility criteria were defined according to the PICO (P, patient; I, intervention; C, comparison; O, outcome)^142^. In brief, we included articles characterizing age-related alterations in articular cartilage and/or subchondral bone of murine knee joints to traumatic injury (i.e., each study had to include both aged and young mice or middle-aged and young mice). Young, middle-aged, and aged mice were defined as 2-7, 9-15, and 18-24 months old, respectively, and correspond to ages 20-30, 38-47, and 56-69 years in humans, respectively^143^. Since middle-aged and aged mice display similar proteomic signatures^112^, this study combined middle-aged and aged mice and defined them as the “aged” group. No restrictions were set according to mouse strain or article publication year. The following articles were excluded: (1) studies that included genetically modified animals, as such models likely oversimplify the disease process, whereas naturally occurring OA is almost certainly polygenic in nature^144^; (2) studies that did not explicitly compare articular cartilage or subchondral bone in the knee joint between young and aged mice; and (3) treatment arm includes another intervention (e.g., saline intra-articular injection), which may interact with aging effects on the disease progression.

Outcome measurements related to assessment of articular cartilage damage or subchondral bone alterations were divided into six categories based on a slightly modified version of previous meta-analyses^145,146^. We used the following hierarchy of outcomes:

1. morphology (e.g., Osteoarthritis Research Society International [OARSI] score^147^, Mankin’s score^148^, MRI based classifications of morphological changes);
2. morphometry (any kind of quantitative methods on microscopic images of articular cartilage and subchondral bone including computational image analysis techniques^149^);
3. composition (any kind of quantitative methods for quantification of proteoglycan or collagen);
4. biomechanical characterization (e.g., tensile and compressive measures of stiffness);
5. biomarker (e.g., mirco RNA in synovial fluid);
6. molecular biology (e.g., ECM-related gene expression and protein synthesis)

PubMed was used for the electronic database search. Google Scholar was also used as a complementary search engine. A manual search of the reference lists of past reviews was performed^150^. Furthermore, citation searching was performed on the original records using Web of Science. These citation indexes are recommended by the Cochrane Handbook^140^. For the database search was performed on November 14, 2022. Electronic searches for PubMed used combined key terms, including “Animal, Laboratory” “Aging,” “Age factors,” “Cartilage, Articular,” and “Osteoarthritis” using Medical Subject Headings terms.

A single reviewer (HI), as a content and statistical expert, assessed eligibility in accordance with the Cochran Handbook recommendations^140^. The reviewer screened titles and abstracts yielded by the search. Full manuscripts of the articles that met the eligibility criteria were then obtained and reviewed. During these processes, the reviewer prepared and used simple, pre-designed Google spreadsheets to assess eligibility by extracting study features.

The same reviewer extracted data regarding basic study information (authors, publication year, and country of corresponding author), experimental condition (i.e., mice strain, age, sample size, and sex), target joint (tibiofemoral or patellofemoral joints), and outcome measures. If outcome measures from multiple time points were reported within the same age category (e.g., three and six months old from the young group), we averaged the effect size^151^. If outcome measures from multiple compartments (e.g., medial and lateral compartments in tibiofemoral joint and patellofemoral joint) were presented, data from the most severe region were extracted. When data were not reported or were unclear, we contacted the authors directly. A reminder was sent to those who had not replied. If data were provided only in figures, the graphically presented data was converted to numerical data using a reliable and validated digital ruler software (WebPlotDigitizer)^152,153^. We have previously confirmed the excellent inter-rater reliability between two independent reviewers (intraclass correlation coefficient [2,1]: 0.999)^154^.

#### Determining genes upregulated in murine knee joint after traumatic injury

We have accessed archived RNA-seq data from young (3 months) and aged (15 months) mice from the study published by Sebastian et al^26^, which yielded 2,738 genes identified across the different time points (day1, week1, week2, and week6) after ACL rupture. We then isolated differentially expressed (false discovery rate <0.05) genes from young+ACL rupture at week1 (935 genes), young+ACL rupture at week2 (1,121 genes), aged+ACL rupture at week1 (1,422 genes), and aged+ACL rupture at week2 (1,461 genes). Using VENN DIAGRAMS software (developed by Van de Peer Lab; http://bioinformatics.psb.ugent.be/webtools/Venn/), we finally identified 241 genes upregulated uniquely in aged murine knee joint after ACL rupture (see **Table S2**). Further, we also identified 726 genes that were upregulated after ACL rupture in both young and aged murine knee joints. We then used these gene lists in the subsequent analyses for the tissue-specific network construction shown below.

We have also repeated the same procedure for the different set of transcriptomic data of aged murine knee joint after DMM induction^8^. We have accessed archived microarray data from young (3 months) and aged (12 months) mice from the study published by Loeser RF^8^, which yielded differentially expressed 549 probe sets with 432 genes in young or aged murine knee joint at 8 weeks after DMM induction. We then isolated 251 probe sets with 203 genes upregulated uniquely in aged murine knee joint at 8 weeks after DMM induction and used for subsequent analyses.

#### Determining genes associated with increased ECM stiffness of articular cartilage in murine knee joint

We sought to define genes associated with ECM stiffness in cartilage via a systematic literature search in accordance with the guidelines shown above^137-141^. We included articles investigating the effects of loss- or gain-of-function approaches on ECM stiffness in cartilage of murine knee joint. No restrictions were set according to mouse age, strain, or article publication year. PubMed was used for electronic database search. Google Scholar was also used as a complementary search engine. A manual search of the reference lists of past reviews was performed^155^. Furthermore, citation searching was performed on the original records using Web of Science as recommended^140^. The database search was performed on November 15, 2022. Electronic searches for PubMed used combined key terms, including “Animal, Genetically Modified” “Cartilage, Articular,” and “Osteoarthritis” using Medical Subject Headings terms.

A single reviewer (HI), as a content and statistical expert, assessed eligibility in accordance with the Cochran Handbook recommendations^140^. The reviewer screened titles and abstracts yielded by the search. Full manuscripts of the articles that met the eligibility criteria were then obtained and reviewed. During these processes, the reviewer prepared and used simple, pre-designed Google spreadsheets to assess eligibility by extracting study features.

When the loss-of- or gain-of-function approach significantly (p <0.05) changed ECM stiffness of articular cartilage evaluated by atomic force microscopy or similar assessment systems designed for assessing biophysical properties of cartilage, we classified the function of genes as (“+”) or (“-”) respectively; and as no effect (“=”) when no statistically significant difference was reported between the mice with the loss-of- or gain-of-function approach and wild type control groups. More specifically, we categorized genes as (“+”) when (1) loss-of-function approach significantly decreased ECM stiffness or (2) gain-of-function approach significantly increased ECM stiffness.

#### Determining articular chondrocyte markers in murine knee joint

Articular chondrocyte markers were defined using previously published single cell RNA-seq data obtained from articular cartilage from 10-week-old adult male murine joint (n = 5)^40^. After excluding immune and blood cells (CD45-positive or TER119-positive cells), the original study has identified 10 different clusters with distinct gene expression profiles, in which five clusters were annotated as “chondrocytes”^40^. We defined chondrocyte markers as the genes which showed significantly higher enrichment to those five chondrocyte clusters compared to the other five clusters, resulting in 42 different genes (see **Table S3**).

#### Identification of cellular origin of cytokines using Tabula Muris Senis

We determined cell type and state identity by leveraging annotations provided in the *Tabula Muris Senis* following the “cell ontology” structure^156^. The *Tabula Muris Senis* is a comprehensive resource for the cell biology community which offers a detailed molecular and cell-type specific portrait of aging^156^. We have used the data provided by droplet experiments. No restrictions were set according to mouse age, sex, and tissue type.

#### Construction of cytokine synergistic network regulating ECM homeostasis

Using systematic literature approach, we have manually curated pair of cytokines that display synergistic impact on ECM remodeling (i.e., MMPs and/or TIMPs) in articular cartilage. For the cytokine of interest, we have focused on cytokines that display synergistic effect with OSM and/or IL6. PubMed was used for electronic database search which was performed on January 14, 2023. Google Scholar was also used as a complementary search engine. Electronic searches for PubMed used combined key terms, including “Chondrocytes,” “Cartilage,” “Interleukin-6,” “Oncostatin M,” and “Drug Synergism” using Medical Subject Headings terms.

A single reviewer (HI), as a content and statistical expert, assessed eligibility. The reviewer screened titles and abstracts yielded by the search. Full manuscripts of the articles that met the eligibility criteria were then obtained and reviewed. During these processes, the reviewer prepared and used simple, pre-designed Google spreadsheets to assess eligibility by extracting study features.

When the combination of two cytokines significantly (p <0.05) changed ECM-related genes or proteins in articular chondrocytes compared to single treatment alone, we classified the pair of cytokines as increased (“+”) or decreased (“-”), or no significant effect (“=”).

#### Pharmacological validation using microarray

We accessed the archived microarray data collected at 4 hours after OSM or IL6 supplementation (100ng/mL) to murine articular chondrocyte^102^. The bioinformatics analysis conducted by the authors, Liu X et al.^102^, revealed 2,373 and 961 differentially (false discovery rate <0.05) expressed genes after OSM and IL6 treatments, respectively. We then used these gene lists in the subsequent analyses for the tissue-specific network construction shown below. To prioritize genes which displayed drastic changes by OSM supplementation, we isolated nine genes significantly upregulated (mean expression >10, log fold change >2.5).

#### Defining a set of genes of “inflammation” phenotype of people with KOA

We have defined the genes associated with “inflammation” phenotype of people with KOA in accordance with single-cell RNA-seq data from KOA cartilage samples (n = 131; mean age: 66.6 years old; 61.8%females; mean joint space narrowing score: 3.91)^105^. Using an unsupervised clustering approach, the cartilage samples were categorized into 1 (n = 81; 61.8%), 2 (n = 24; 18.3%), 3 (n = 10; 7.6%), and 4 (n = 16; 12.2%) according to the top 4000 most variable genes^105^. People classified into the inflammation phenotype (mean age: 71.0 years old; 56.3%females; median Kellgren and Lawrence grade 4) displayed more severe phenotype, as evidenced by severe joint space narrowing (mean joint space narrowing score: 4.38)^105^.

### QUANTIFICATION AND STATISTICAL ANALYSIS

#### Meta-analysis

To characterize age-related alterations in articular cartilage and/or subchondral bone of murine knee joints to traumatic injury, we calculated pooled estimates and 95% confidence intervals for standardized mean differences (SMD) of outcome measures using the DerSimonian-Laird method^157^. This method considers the precision of individual studies and the variation between studies and weighs each study accordingly. SMD were calculated using the mean between-group difference (young vs. young+injury, aged vs. aged+injury) divided by the pooled standard deviation^157^. Meta-analyses were performed using RevMan version 5.3 (Nordic Cochrane Centre, Cochrane Collaboration, Copenhagen, Denmark). Study heterogeneity, defined as the inter-trial variation in study outcomes, was assessed using *I^2^*, which is the proportion of total variance explained by inter-trial heterogeneity^158^. To standardize semi-quantitative scores of cartilage degeneration, all histological scores provided in each included study were converted to 0-100 and recalculated as in a previous meta-analysis^159^, with higher score indicates severe cartilage degeneration.

#### Construction of tissue-specific network

Tissue-specific networks were constructed using HumanBase web tool^33^ with the genes of interest used an input. The data of tissue-specific network data was based on several data types that constituted the underlying network, including experimentally produced protein-protein interactions (http://hb.flatironinstitute.org/data), and the interaction confidence is the edge weight assigned from the algorithm used to create this compendium network. Since the tissue-specific network was established based on human tissue, all the gene symbols were translated into human gene symbols prior to the analysis.

#### Functional characterization of transcriptome using GO enrichment analysis

To determine the biological function of genes of interests, GO enrichment analysis (biological process) was performed by Enrichr software^37^. REVIGO software^133^ was applied to summarize redundant GO terms and visualize the summarized results.

#### GSEA

ssGSEA was performed using GSEA web tool provided by Broad Institute Website (https://www.gsea-msigdb.org/gsea/index.jsp) with gene scores defined by log2 fold change of gene expression profiles of aged+ACL rupture at week1 and 2. As a gene set, we have used genes associated with increased ECM stiffness in cartilage we have defined above.

#### Network propagation using RWR on cartilage-specific global network

On cartilage-specific global network, RWR was performed by R/Bioconductor package RandomWalkRestartMH^98^ with the *Osmr* gene used as a seeded node. After iteratively reaching stability, the affinity score of all nodes in the given network to *Osmr* node were obtained. Cartilage-specific global network was constructed using a gold standard data set (i.e., already known gene interactions) downloaded from HumanBase software (https://hb.flatironinstitute.org/download)^33^. In the statistical analysis that characterize the relationship between the pseudo-activated genes (affinity score >0) and interest of phenotype, we have excluded the 1,284 genes that were significantly changed after traumatic injury in young mice to dissect the effects of age-specific transcriptomic response (i.e., OSM-OSMR axis) by removing the injury effects.

#### Unsupervised machine learning

PCA was performed for data reduction to identify the principal components that represent differences in the given data sets using JMP Pro 16 software (SAS Institute, Cary, NC). PCA produces linear combinations of the original variables to generate the principal components^160^, and visualization was generated by projecting the original data to the first two principal components.

#### Cytokine inference

To define enriched cytokine signaling signatures of given gene expression data, cytokine inference was performed using CytoSig software^71^. CytoSig analyzes defined cytokine signatures that are differentially expressed when a cell is exposed to a specific cytokine (that is name giving for the respective cytokine signature). This study used the data of transcriptomic response (i.e., log2 fold change) of 58 genes related to ECM remodeling across the two groups (young+ACL rupture vs. aged+ACL rupture) over time was used as an input.

#### WGCNA

This study used the WGCNA package to build a weighted gene co-expression network using the archived RNA-seq database (GSE114007)^99^, which finally yielded 13,729 genes identified across the different data sets after filtering low expression genes. We used filterByExpr function for count data in RNA-seq data^161^. The key parameter, β, for weighted network construction was optimized to maintain both the scale-free topology and sufficient node connectivity as recommended in the manual. A topological overlap matrix (TOM) was then formulated based on the adjacency values to calculate the corresponding dissimilarity (1-TOM) values. Module identification was accomplished with the dynamic tree cut method by hierarchically clustering genes using 1-TOM as the distance measure with a minimum size cutoff of 30 and a deep split value of 2 for the resulting dendrogram. A module preservation function was used to verify the stability of the identified modules by calculating module preservation and quality statistics in the WGCNA package^101^.

#### Drug repurposing via ConnectivityMap analysis

We explored chemical compounds that could possibly reverse OSM signaling, injured aged knees, and inflammation phenotype in KOA using ConnectivityMap^106^. Using the online interface clue.io (https://clue.io/), we submitted the genes with OSM signaling pathway, age-related stress response, and inflammation phenotype to calculate a ‘tau’ connectivity score to gene expression signatures experimentally induced by various perturbations in nine cell lines. Gene list for OSM signaling pathway was obtained from “WikiPathway_2021_Human” via Enrichr (https://maayanlab.cloud/Enrichr/). Since all the included genes across three conditions (i.e., OSM signaling, age-related stress response, and inflammation phenotype) were upregulated genes, the subsequent analysis has focused on a negative tau score (gene expression signature of a perturbation opposes the submitted query). Recommended thresholds for further consideration of results are tau of below −90 (https://clue.io/connectopedia/connectivity_scores). Of 2837 compounds were evaluated in clue.io, we shortlisted compounds where the summary tau across cell lines were lower than −90.

#### Statistical analysis

All statistical analyses were performed using JMP Pro 16 software (SAS Institute, Cary, NC) or SPSS Statistics for Windows, Version 28.0 (IBM Corp., NY, USA). Except where indicated, data are displayed as means, with uncertainty expressed as 95% confidence intervals (mean ± 95% CI). Two-tailed Student *t*-test, linear regression analysis, logistic regression analysis, or Fisher’s exact test were used for statistical analyses. We checked the features of the regression model by comparing the residuals vs. fitted values (i.e., the residuals had to be normally distributed around zero) and independence between observations. No correction was applied for multiple comparison because outcomes were determined a priori and were highly correlated. No statistical analyses included confounders (e.g., body mass in each animal) due to the small sample size. We conducted a complete-case analysis in the case of missing data. In all experiments, p-values <0.05 were considered statistically significant. Throughout this text, “*n*” represents the number of independent observations of knees or cells from different animals. Specific data representation details and statistical procedures are also indicated in the figure legends.

## References

1. López-Otín, C., Blasco, M.A., Partridge, L., Serrano, M., and Kroemer, G. (2013). The hallmarks of aging. Cell 153, 1194–1217. 10.1016/j.cell.2013.05.039.

2. Kennedy, B.K., Berger, S.L., Brunet, A., Campisi, J., Cuervo, A.M., Epel, E.S., Franceschi, C., Lithgow, G.J., Morimoto, R.I., Pessin, J.E., et al. (2014). Geroscience: linking aging to chronic disease. Cell 159, 709–713. 10.1016/j.cell.2014.10.039.

3. Loeser, R.F., Collins, J.A., and Diekman, B.O. (2016). Ageing and the pathogenesis of osteoarthritis. Nat Rev Rheumatol 12, 412–420. 10.1038/nrrheum.2016.65.

4. Guilak, F. (2011). Biomechanical factors in osteoarthritis. Best Pract Res Clin Rheumatol 25, 815–823. 10.1016/j.berh.2011.11.013.

5. Iijima, H., Gilmer, G., Wang, K., Sivakumar, S., Evans, C., Matsui, Y., and Ambrosio, F. (2022). Meta-analysis Integrated With Multi-omics Data Analysis to Elucidate Pathogenic Mechanisms of Age-Related Knee Osteoarthritis in Mice. J Gerontol A Biol Sci Med Sci 77, 1321–1334. 10.1093/gerona/glab386.

6. Lotz, M., and Loeser, R.F. (2012). Effects of aging on articular cartilage homeostasis. Bone 51, 241–248. 10.1016/j.bone.2012.03.023.

7. Houtman, E., Tuerlings, M., Riechelman, J., Suchiman, E., van der Wal, R.J.P., Nelissen, R., Mei, H., Ramos, Y.F.M., Coutinho de Almeida, R., and Meulenbelt, I. (2021). Elucidating mechano-pathology of osteoarthritis: transcriptome-wide differences in mechanically stressed aged human cartilage explants. Arthritis Res Ther 23, 215. 10.1186/s13075-021-02595-8.

8. Loeser, R.F., Olex, A.L., McNulty, M.A., Carlson, C.S., Callahan, M.F., Ferguson, C.M., Chou, J., Leng, X., and Fetrow, J.S. (2012). Microarray analysis reveals age-related differences in gene expression during the development of osteoarthritis in mice. Arthritis Rheum 64, 705–717. 10.1002/art.33388.

9. Madej, W., van Caam, A., Davidson, E.N., Hannink, G., Buma, P., and van der Kraan, P.M. (2016). Ageing is associated with reduction of mechanically-induced activation of Smad2/3P signaling in articular cartilage. Osteoarthritis Cartilage 24, 146–157. 10.1016/j.joca.2015.07.018.

10. Kurz, B., Jin, M., Patwari, P., Cheng, D.M., Lark, M.W., and Grodzinsky, A.J. (2001). Biosynthetic response and mechanical properties of articular cartilage after injurious compression. J Orthop Res 19, 1140–1146. 10.1016/s0736-0266(01)00033-x.

11. Englund, M., Guermazi, A., Gale, D., Hunter, D.J., Aliabadi, P., Clancy, M., and Felson, D.T. (2008). Incidental meniscal findings on knee MRI in middle-aged and elderly persons. N Engl J Med 359, 1108–1115. 10.1056/NEJMoa0800777.

12. Hasegawa, A., Otsuki, S., Pauli, C., Miyaki, S., Patil, S., Steklov, N., Kinoshita, M., Koziol, J., D’Lima, D.D., and Lotz, M.K. (2012). Anterior cruciate ligament changes in the human knee joint in aging and osteoarthritis. Arthritis Rheum 64, 696–704. 10.1002/art.33417.

13. Loeser, R.F., Goldring, S.R., Scanzello, C.R., and Goldring, M.B. (2012). Osteoarthritis: a disease of the joint as an organ. Arthritis Rheum 64, 1697–1707. 10.1002/art.34453.

14. Chou, C.H., Jain, V., Gibson, J., Attarian, D.E., Haraden, C.A., Yohn, C.B., Laberge, R.M., Gregory, S., and Kraus, V.B. (2020). Synovial cell cross-talk with cartilage plays a major role in the pathogenesis of osteoarthritis. Sci Rep 10, 10868. 10.1038/s41598-020-67730-y.

15. Zhang, F., Wei, K., Slowikowski, K., Fonseka, C.Y., Rao, D.A., Kelly, S., Goodman, S.M., Tabechian, D., Hughes, L.B., Salomon-Escoto, K., et al. (2019). Defining inflammatory cell states in rheumatoid arthritis joint synovial tissues by integrating single-cell transcriptomics and mass cytometry. Nat Immunol 20, 928–942. 10.1038/s41590-019-0378-1.

16. Zhang, F., Jonsson, A.H., Nathan, A., Wei, K., Millard, N., Xiao, Q., Gutierrez-Arcelus, M., Apruzzese, W., Watts, G.F.M., Weisenfeld, D., et al. (2022). Cellular deconstruction of inflamed synovium defines diverse inflammatory phenotypes in rheumatoid arthritis. bioRxiv, 2022.2002.2025.481990. 10.1101/2022.02.25.481990.

17. Kapoor, M., Martel-Pelletier, J., Lajeunesse, D., Pelletier, J.P., and Fahmi, H. (2011). Role of proinflammatory cytokines in the pathophysiology of osteoarthritis. Nat Rev Rheumatol 7, 33–42. 10.1038/nrrheum.2010.196.

18. Barabási, A.L., Gulbahce, N., and Loscalzo, J. (2011). Network medicine: a network-based approach to human disease. Nat Rev Genet 12, 56–68. 10.1038/nrg2918.

19. Wong, A.K., Krishnan, A., and Troyanskaya, O.G. (2018). GIANT 2.0: genome-scale integrated analysis of gene networks in tissues. Nucleic Acids Res 46, W65–w70. 10.1093/nar/gky408.

20. Dell’Isola, A., Allan, R., Smith, S.L., Marreiros, S.S., and Steultjens, M. (2016). Identification of clinical phenotypes in knee osteoarthritis: a systematic review of the literature. BMC Musculoskelet Disord 17, 425. 10.1186/s12891-016-1286-2.

21. Faust, H.J., Zhang, H., Han, J., Wolf, M.T., Jeon, O.H., Sadtler, K., Peña, A.N., Chung, L., Maestas, D.R., and Tam, A.J. (2020). IL-17 and immunologically induced senescence regulate response to injury in osteoarthritis. The Journal of clinical investigation 130, 5493–5507.

22. Huang, H., Veien, E.S., Zhang, H., Ayers, D.C., and Song, J. (2016). Skeletal characterization of Smurf2-deficient mice and in vitro analysis of Smurf2-deficient chondrocytes. PLoS One 11, e0148088.

23. Huang, H., Skelly, J.D., Ayers, D.C., and Song, J. (2017). Age-dependent changes in the articular cartilage and subchondral bone of C57BL/6 mice after surgical destabilization of medial meniscus. Scientific reports 7, 1–9.

24. Ko, F.C., Dragomir, C., Plumb, D.A., Goldring, S.R., Wright, T.M., Goldring, M.B., and van der Meulen, M.C. (2013). In vivo cyclic compression causes cartilage degeneration and subchondral bone changes in mouse tibiae. Arthritis & Rheumatism 65, 1569–1578.

25. Loeser, R.F., Olex, A.L., McNulty, M.A., Carlson, C.S., Callahan, M.F., Ferguson, C.M., Chou, J., Leng, X., and Fetrow, J.S. (2012). Microarray analysis reveals age_related differences in gene expression during the development of osteoarthritis in mice. Arthritis & Rheumatism 64, 705–717.

26. Sebastian, A., Murugesh, D.K., Mendez, M.E., Hum, N.R., Rios-Arce, N.D., McCool, J.L., Christiansen, B.A., and Loots, G.G. (2020). Global Gene Expression Analysis Identifies Age-Related Differences in Knee Joint Transcriptome during the Development of Post-Traumatic Osteoarthritis in Mice. Int J Mol Sci 21. 10.3390/ijms21010364.

27. Kuyinu, E.L., Narayanan, G., Nair, L.S., and Laurencin, C.T. (2016). Animal models of osteoarthritis: classification, update, and measurement of outcomes. J Orthop Surg Res 11, 19. 10.1186/s13018-016-0346-5.

28. Diekman, B.O., Sessions, G.A., Collins, J.A., Knecht, A.K., Strum, S.L., Mitin, N.K., Carlson, C.S., Loeser, R.F., and Sharpless, N.E. (2018). Expression of p16(INK)(4a) is a biomarker of chondrocyte aging but does not cause osteoarthritis. Aging Cell 17, e12771. 10.1111/acel.12771.

29. Faust, H.J., Zhang, H., Han, J., Wolf, M.T., Jeon, O.H., Sadtler, K., Peña, A.N., Chung, L., Maestas, D.R., Jr., Tam, A.J., et al. (2020). IL-17 and immunologically induced senescence regulate response to injury in osteoarthritis. J Clin Invest 130, 5493–5507. 10.1172/jci134091.

30. Iijima, H., Aoyama, T., Ito, A., Tajino, J., Nagai, M., Zhang, X., Yamaguchi, S., Akiyama, H., and Kuroki, H. (2014). Destabilization of the medial meniscus leads to subchondral bone defects and site-specific cartilage degeneration in an experimental rat model. Osteoarthritis Cartilage 22, 1036–1043. 10.1016/j.joca.2014.05.009.

31. Iijima, H., Aoyama, T., Tajino, J., Ito, A., Nagai, M., Yamaguchi, S., Zhang, X., Kiyan, W., and Kuroki, H. (2016). Subchondral plate porosity colocalizes with the point of mechanical load during ambulation in a rat knee model of post-traumatic osteoarthritis. Osteoarthritis Cartilage 24, 354–363. 10.1016/j.joca.2015.09.001.

32. Li, G., Yin, J., Gao, J., Cheng, T.S., Pavlos, N.J., Zhang, C., and Zheng, M.H. (2013). Subchondral bone in osteoarthritis: insight into risk factors and microstructural changes. Arthritis Res Ther 15, 223. 10.1186/ar4405.

33. Greene, C.S., Krishnan, A., Wong, A.K., Ricciotti, E., Zelaya, R.A., Himmelstein, D.S., Zhang, R., Hartmann, B.M., Zaslavsky, E., Sealfon, S.C., et al. (2015). Understanding multicellular function and disease with human tissue-specific networks. Nat Genet 47, 569–576. 10.1038/ng.3259.

34. Lage, K., Hansen, N.T., Karlberg, E.O., Eklund, A.C., Roque, F.S., Donahoe, P.K., Szallasi, Z., Jensen, T.S., and Brunak, S. (2008). A large-scale analysis of tissue-specific pathology and gene expression of human disease genes and complexes. Proc Natl Acad Sci U S A 105, 20870–20875. 10.1073/pnas.0810772105.

35. Reverter, A., Ingham, A., and Dalrymple, B.P. (2008). Mining tissue specificity, gene connectivity and disease association to reveal a set of genes that modify the action of disease causing genes. BioData Min 1, 8. 10.1186/1756-0381-1-8.

36. Baker, B.M., Trappmann, B., Wang, W.Y., Sakar, M.S., Kim, I.L., Shenoy, V.B., Burdick, J.A., and Chen, C.S. (2015). Cell-mediated fibre recruitment drives extracellular matrix mechanosensing in engineered fibrillar microenvironments. Nat Mater 14, 1262–1268. 10.1038/nmat4444.

37. Kuleshov, M.V., Jones, M.R., Rouillard, A.D., Fernandez, N.F., Duan, Q., Wang, Z., Koplev, S., Jenkins, S.L., Jagodnik, K.M., Lachmann, A., et al. (2016). Enrichr: a comprehensive gene set enrichment analysis web server 2016 update. Nucleic Acids Res 44, W90–97. 10.1093/nar/gkw377.

38. Kagan, H.M., and Li, W. (2003). Lysyl oxidase: properties, specificity, and biological roles inside and outside of the cell. J Cell Biochem 88, 660–672. 10.1002/jcb.10413.

39. Kim, J.H., Lee, G., Won, Y., Lee, M., Kwak, J.S., Chun, C.H., and Chun, J.S. (2015). Matrix cross-linking-mediated mechanotransduction promotes posttraumatic osteoarthritis. Proc Natl Acad Sci U S A 112, 9424–9429. 10.1073/pnas.1505700112.

40. Sebastian, A., McCool, J.L., Hum, N.R., Murugesh, D.K., Wilson, S.P., Christiansen, B.A., and Loots, G.G. (2021). Single-Cell RNA-Seq Reveals Transcriptomic Heterogeneity and Post-Traumatic Osteoarthritis-Associated Early Molecular Changes in Mouse Articular Chondrocytes. Cells 10. 10.3390/cells10061462.

41. Stolz, M., Gottardi, R., Raiteri, R., Miot, S., Martin, I., Imer, R., Staufer, U., Raducanu, A., Düggelin, M., Baschong, W., et al. (2009). Early detection of aging cartilage and osteoarthritis in mice and patient samples using atomic force microscopy. Nat Nanotechnol 4, 186–192. 10.1038/nnano.2008.410.

42. Schmauck-Medina, T., Molière, A., Lautrup, S., Zhang, J., Chlopicki, S., Madsen, H.B., Cao, S., Soendenbroe, C., Mansell, E., Vestergaard, M.B., et al. (2022). New hallmarks of ageing: a 2022 Copenhagen ageing meeting summary. Aging (Albany NY) 14, 6829–6839. 10.18632/aging.204248.

43. Zhong, W., Li, Y., Li, L., Zhang, W., Wang, S., and Zheng, X. (2013). YAP-mediated regulation of the chondrogenic phenotype in response to matrix elasticity. J Mol Histol 44, 587–595. 10.1007/s10735-013-9502-y.

44. Du, J., Zu, Y., Li, J., Du, S., Xu, Y., Zhang, L., Jiang, L., Wang, Z., Chien, S., and Yang, C. (2016). Extracellular matrix stiffness dictates Wnt expression through integrin pathway. Sci Rep 6, 20395. 10.1038/srep20395.

45. Iijima, H., Gilmer, G., Wang, K., Bean, A.C., He, Y., Lin, H., Tang, W.-Y., Lamont, D., Tai, C., Ito, A., et al. (2023). Age-related matrix stiffening epigenetically regulates α-Klotho expression and compromises chondrocyte integrity. Nature Communications 14, 18. 10.1038/s41467-022-35359-2.

46. Alberton, P., Dugonitsch, H.C., Hartmann, B., Li, P., Farkas, Z., Saller, M.M., Clausen-Schaumann, H., and Aszodi, A. (2019). Aggrecan hypomorphism compromises articular cartilage biomechanical properties and is associated with increased incidence of spontaneous osteoarthritis. International Journal of Molecular Sciences 20, 1008.

47. Alexopoulos, L.G., Youn, I., Bonaldo, P., and Guilak, F. (2009). Developmental and osteoarthritic changes in Col6a1_knockout mice: biomechanics of type VI collagen in the cartilage pericellular matrix. Arthritis & Rheumatism: Official Journal of the American College of Rheumatology 60, 771–779.

48. Batista, M.A., Nia, H.T., Önnerfjord, P., Cox, K.A., Ortiz, C., Grodzinsky, A.J., Heinegård, D., and Han, L. (2014). Nanomechanical phenotype of chondroadherin-null murine articular cartilage. Matrix Biology 38, 84–90.

49. Candela, M.E., Wang, C., Gunawardena, A.T., Zhang, K., Cantley, L., Yasuhara, R., Usami, Y., Francois, N., Iwamoto, M., and van der Flier, A. (2016). Alpha 5 integrin mediates osteoarthritic changes in mouse knee joints. PloS one 11, e0156783.

50. Chen, Y., Cossman, J., Jayasuriya, C.T., Li, X., Guan, Y., Fonseca, V., Yang, K., Charbonneau, C., Yu, H., and Kanbe, K. (2016). Deficient mechanical activation of anabolic transcripts and post-traumatic cartilage degeneration in matrilin-1 knockout mice. PLoS One 11, e0156676.

51. Christensen, S.E., Coles, J.M., Zelenski, N.A., Furman, B.D., Leddy, H.A., Zauscher, S., Bonaldo, P., and Guilak, F. (2012). Altered trabecular bone structure and delayed cartilage degeneration in the knees of collagen VI null mice. PloS one 7, e33397.

52. Coles, J.M., Zhang, L., Blum, J.J., Warman, M.L., Jay, G.D., Guilak, F., and Zauscher, S. (2010). Loss of cartilage structure, stiffness, and frictional properties in mice lacking PRG4. Arthritis & Rheumatism 62, 1666–1674.

53. Gamer, L.W., Pregizer, S., Gamer, J., Feigenson, M., Ionescu, A., Li, Q., Han, L., and Rosen, V. (2018). The role of Bmp2 in the maturation and maintenance of the murine knee joint. Journal of Bone and Mineral Research 33, 1708–1717.

54. Gronau, T., Krüger, K., Prein, C., Aszodi, A., Gronau, I., Iozzo, R.V., Mooren, F.C., Clausen-Schaumann, H., Bertrand, J., and Pap, T. (2017). Forced exercise-induced osteoarthritis is attenuated in mice lacking the small leucine-rich proteoglycan decorin. Annals of the Rheumatic Diseases 76, 442–449.

55. Han, B., Li, Q., Wang, C., Patel, P., Adams, S.M., Doyran, B., Nia, H.T., Oftadeh, R., Zhou, S., and Li, C.Y. (2019). Decorin regulates the aggrecan network integrity and biomechanical functions of cartilage extracellular matrix. ACS nano 13, 11320–11333.

56. Han, B., Li, Q., Wang, C., Chandrasekaran, P., Zhou, Y., Qin, L., Liu, X.S., Enomoto-Iwamoto, M., Kong, D., and Iozzo, R.V. (2021). Differentiated activities of decorin and biglycan in the progression of post-traumatic osteoarthritis. Osteoarthritis and Cartilage 29, 1181–1192.

57. Jia, H., Ma, X., Tong, W., Doyran, B., Sun, Z., Wang, L., Zhang, X., Zhou, Y., Badar, F., and Chandra, A. (2016). EGFR signaling is critical for maintaining the superficial layer of articular cartilage and preventing osteoarthritis initiation. Proceedings of the National Academy of Sciences 113, 14360–14365.

58. Li, P., Fleischhauer, L., Nicolae, C., Prein, C., Farkas, Z., Saller, M.M., Prall, W.C., Wagener, R., Heilig, J., and Niehoff, A. (2020). Mice lacking the matrilin family of extracellular matrix proteins develop mild skeletal abnormalities and are susceptible to age-associated osteoarthritis. International journal of molecular sciences 21, 666.

59. Li, Q., Han, B., Wang, C., Tong, W., Wei, Y., Tseng, W.J., Han, L.H., Liu, X.S., Enomoto_Iwamoto, M., and Mauck, R.L. (2020). Mediation of cartilage matrix degeneration and fibrillation by decorin in post_traumatic osteoarthritis. Arthritis & Rheumatology 72, 1266–1277.

60. Stolz, M., Gottardi, R., Raiteri, R., Miot, S., Martin, I., Imer, R., Staufer, U., Raducanu, A., Düggelin, M., and Baschong, W. (2009). Early detection of aging cartilage and osteoarthritis in mice and patient samples using atomic force microscopy. Nature nanotechnology 4, 186–192.

61. Wang, C., Brisson, B.K., Terajima, M., Li, Q., Han, B., Goldberg, A.M., Liu, X.S., Marcolongo, M.S., Enomoto-Iwamoto, M., and Yamauchi, M. (2020). Type III collagen is a key regulator of the collagen fibrillar structure and biomechanics of articular cartilage and meniscus. Matrix Biology 85, 47–67.

62. Wei, Y., Ma, X., Sun, H., Gui, T., Li, J., Yao, L., Zhong, L., Yu, W., Han, B., and Nelson, C.L. (2022). EGFR signaling is required for maintaining adult cartilage homeostasis and attenuating osteoarthritis progression. Journal of Bone and Mineral Research 37, 1012–1023.

63. Xu, L., Flahiff, C., Waldman, B., Wu, D., Olsen, B., Setton, L., and Li, Y. (2003). Osteoarthritis_like changes and decreased mechanical function of articular cartilage in the joints of mice with the chondrodysplasia gene (cho). Arthritis & Rheumatism: Official Journal of the American College of Rheumatology 48, 2509–2518.

64. Xu, X., Li, Z., Leng, Y., Neu, C.P., and Calve, S. (2016). Knockdown of the pericellular matrix molecule perlecan lowers in situ cell and matrix stiffness in developing cartilage. Developmental biology 418, 242–247.

65. Szklarczyk, D., Kirsch, R., Koutrouli, M., Nastou, K., Mehryary, F., Hachilif, R., Gable, A.L., Fang, T., Doncheva, N.T., Pyysalo, S., et al. (2022). The STRING database in 2023: protein-protein association networks and functional enrichment analyses for any sequenced genome of interest. Nucleic Acids Res. 10.1093/nar/gkac1000.

66. Boer, C.G., Hatzikotoulas, K., Southam, L., Stefánsdóttir, L., Zhang, Y., de Almeida, R.C., Wu, T.T., Zheng, J., Hartley, A., and Teder-Laving, M. (2021). Deciphering osteoarthritis genetics across 826,690 individuals from 9 populations. Cell 184, 4784–4818. e4717.

67. Rodriguez, R., Seegmiller, R., Stark, M., and Bridgewater, L. (2004). A type XI collagen mutation leads to increased degradation of type II collagen in articular cartilage. Osteoarthritis and cartilage 12, 314–320.

68. Barbie, D.A., Tamayo, P., Boehm, J.S., Kim, S.Y., Moody, S.E., Dunn, I.F., Schinzel, A.C., Sandy, P., Meylan, E., Scholl, C., et al. (2009). Systematic RNA interference reveals that oncogenic KRAS-driven cancers require TBK1. Nature 462, 108–112. 10.1038/nature08460.

69. Wynn, T.A., and Ramalingam, T.R. (2012). Mechanisms of fibrosis: therapeutic translation for fibrotic disease. Nat Med 18, 1028–1040. 10.1038/nm.2807.

70. Lieberthal, J., Sambamurthy, N., and Scanzello, C.R. (2015). Inflammation in joint injury and post-traumatic osteoarthritis. Osteoarthritis and cartilage 23, 1825–1834.

71. Jiang, P., Zhang, Y., Ru, B., Yang, Y., Vu, T., Paul, R., Mirza, A., Altan-Bonnet, G., Liu, L., Ruppin, E., et al. (2021). Systematic investigation of cytokine signaling activity at the tissue and single-cell levels. Nat Methods 18, 1181–1191. 10.1038/s41592-021-01274-5.

72. Rose-John, S. (2018). Interleukin-6 family cytokines. Cold Spring Harbor perspectives in biology 10, a028415.

73. Liao, Y., Ren, Y., Luo, X., Mirando, A.J., Long, J.T., Leinroth, A., Ji, R.-R., and Hilton, M.J. (2022). Interleukin-6 signaling mediates cartilage degradation and pain in posttraumatic osteoarthritis in a sex-specific manner. Science signaling 15, eabn7082.

74. Richette, P., Latourte, A., Sellam, J., Wendling, D., Piperno, M., Goupille, P., Pers, Y.M., Eymard, F., Ottaviani, S., Ornetti, P., et al. (2021). Efficacy of tocilizumab in patients with hand osteoarthritis: double blind, randomised, placebo-controlled, multicentre trial. Ann Rheum Dis 80, 349–355. 10.1136/annrheumdis-2020-218547.

75. Richards, C.D. (2013). The enigmatic cytokine oncostatin m and roles in disease. ISRN Inflamm 2013, 512103. 10.1155/2013/512103.

76. Barksby, H., Hui, W., Wappler, I., Peters, H., Milner, J., Richards, C., Cawston, T., and Rowan, A. (2006). Interleukin_1 in combination with oncostatin M up_regulates multiple genes in chondrocytes: implications for cartilage destruction and repair. Arthritis & Rheumatism: Official Journal of the American College of Rheumatology 54, 540–550.

77. Catterall, J., Carrere, S., Koshy, P., Degnan, B., Shingleton, W., Brinckerhoff, C., Rutter, J., Cawston, T., and Rowan, A. (2001). Synergistic induction of matrix metalloproteinase 1 by interleukin_1α and oncostatin M in human chondrocytes involves signal transducer and activator of transcription and activator protein 1 transcription factors via a novel mechanism. Arthritis & Rheumatism 44, 2296–2310.

78. Cawston, T., Billington, C., Cleaver, C., Elliott, S., Hui, W., Koshy, P., Shingleton, B., and Rowan, A. (1999). The regulation of MMPs and TIMPs in cartilage turnover. Annals of the New York Academy of Sciences 878, 120–129.

79. Cawston, T., Curry, V., Summers, C., Clark, I., Riley, G., Life, P., Spaull, J., Goldring, M., Koshy, P., and Rowan, A. (1998). The role of oncostatin M in animal and human connective tissue collagen turnover and its localization within the rheumatoid joint. Arthritis & Rheumatism: Official Journal of the American College of Rheumatology 41, 1760–1771.

80. Hui, W., Rowan, A., Richards, C., and Cawston, T. (2003). Oncostatin M in combination with tumor necrosis factor α induces cartilage damage and matrix metalloproteinase expression in vitro and in vivo. Arthritis & Rheumatism: Official Journal of the American College of Rheumatology 48, 3404–3418.

81. Koshy, P., Henderson, N., Logan, C., Life, P., Cawston, T., and Rowan, A. (2002). Interleukin 17 induces cartilage collagen breakdown: novel synergistic effects in combination with proinflammatory cytokines. Annals of the rheumatic diseases 61, 704–713.

82. Koshy, P., Lundy, C., Rowan, A., Porter, S., Edwards, D., Hogan, A., Clark, I., and Cawston, T. (2002). The modulation of matrix metalloproteinase and ADAM gene expression in human chondrocytes by interleukin_1 and oncostatin M: a time_course study using real_time quantitative reverse transcription–polymerase chain reaction. Arthritis & Rheumatism: Official Journal of the American College of Rheumatology 46, 961–967.

83. Litherland, G.J., Dixon, C., Lakey, R.L., Robson, T., Jones, D., Young, D.A., Cawston, T.E., and Rowan, A.D. (2008). Synergistic collagenase expression and cartilage collagenolysis are phosphatidylinositol 3-kinase/Akt signaling-dependent. Journal of Biological Chemistry 283, 14221–14229.

84. Moran, E.M., Mullan, R., McCormick, J., Connolly, M., Sullivan, O., FitzGerald, O., Bresnihan, B., Veale, D.J., and Fearon, U. (2009). Human rheumatoid arthritis tissue production of IL-17A drives matrix and cartilage degradation: synergy with tumour necrosis factor-α, Oncostatin M and response to biologic therapies. Arthritis research & therapy 11, 1–12.

85. Rowan, A.D., Hui, W., Cawston, T.E., and Richards, C.D. (2003). Adenoviral gene transfer of interleukin-1 in combination with oncostatin M induces significant joint damage in a murine model. The American journal of pathology 162, 1975–1984.

86. Rowan, A., Koshy, P., Shingleton, W., Degnan, B., Heath, J., Vernallis, A., Spaull, J., Life, P., Hudson, K., and Cawston, T. (2001). Synergistic effects of glycoprotein 130 binding cytokines in combination with interleukin_1 on cartilage collagen breakdown. Arthritis & Rheumatism: Official Journal of the American College of Rheumatology 44, 1620–1632.

87. Sanchez, C., Deberg, M.A., Burton, S., Devel, P., Reginster, J.-Y.L., and Henrotin, Y.E. (2004). Differential regulation of chondrocyte metabolism by oncostatin M and interleukin-6. Osteoarthritis and cartilage 12, 801–810.

88. Stawski, L., and Trojanowska, M. (2019). Oncostatin M and its role in fibrosis. Connect Tissue Res 60, 40–49. 10.1080/03008207.2018.1500558.

89. Beekhuizen, M., van Osch, G.J., Bot, A.G., Hoekstra, M.C., Saris, D.B., Dhert, W.J., and Creemers, L.B. (2013). Inhibition of oncostatin M in osteoarthritic synovial fluid enhances GAG production in osteoarthritic cartilage repair. Eur Cell Mater 26, 80–90; discussion 90. 10.22203/ecm.v026a06.

90. Tsuchida, A.I., Beekhuizen, M., t Hart, M.C., Radstake, T.R., Dhert, W.J., Saris, D.B., van Osch, G.J., and Creemers, L.B. (2014). Cytokine profiles in the joint depend on pathology, but are different between synovial fluid, cartilage tissue and cultured chondrocytes. Arthritis Res Ther 16, 441. 10.1186/s13075-014-0441-0.

91. Jin, S., Guerrero-Juarez, C.F., Zhang, L., Chang, I., Ramos, R., Kuan, C.H., Myung, P., Plikus, M.V., and Nie, Q. (2021). Inference and analysis of cell-cell communication using CellChat. Nat Commun 12, 1088. 10.1038/s41467-021-21246-9.

92. Single-cell transcriptomics of 20 mouse organs creates a Tabula Muris. (2018). Nature 562, 367-372. 10.1038/s41586-018-0590-4.

93. Li, M., Yin, H., Yan, Z., Li, H., Wu, J., Wang, Y., Wei, F., Tian, G., Ning, C., Li, H., et al. (2022). The immune microenvironment in cartilage injury and repair. Acta Biomater 140, 23–42. 10.1016/j.actbio.2021.12.006.

94. Haraden, C.A., Huebner, J.L., Hsueh, M.F., Li, Y.J., and Kraus, V.B. (2019). Synovial fluid biomarkers associated with osteoarthritis severity reflect macrophage and neutrophil related inflammation. Arthritis Res Ther 21, 146. 10.1186/s13075-019-1923-x.

95. Manukyan, G., Gallo, J., Mikulkova, Z., Trajerova, M., Savara, J., Slobodova, Z., Fidler, E., Shrestha, B., and Kriegova, E. (2023). Phenotypic and functional characterisation of synovial fluid-derived neutrophils in knee osteoarthritis and knee infection. Osteoarthritis Cartilage 31, 72–82. 10.1016/j.joca.2022.09.011.

96. Laan, M., Cui, Z.H., Hoshino, H., Lötvall, J., Sjöstrand, M., Gruenert, D.C., Skoogh, B.E., and Lindén, A. (1999). Neutrophil recruitment by human IL-17 via C-X-C chemokine release in the airways. J Immunol 162, 2347–2352.

97. Cowen, L., Ideker, T., Raphael, B.J., and Sharan, R. (2017). Network propagation: a universal amplifier of genetic associations. Nat Rev Genet 18, 551–562. 10.1038/nrg.2017.38.

98. Valdeolivas, A., Tichit, L., Navarro, C., Perrin, S., Odelin, G., Levy, N., Cau, P., Remy, E., and Baudot, A. (2019). Random walk with restart on multiplex and heterogeneous biological networks. Bioinformatics 35, 497–505. 10.1093/bioinformatics/bty637.

99. Fisch, K.M., Gamini, R., Alvarez-Garcia, O., Akagi, R., Saito, M., Muramatsu, Y., Sasho, T., Koziol, J.A., Su, A.I., and Lotz, M.K. (2018). Identification of transcription factors responsible for dysregulated networks in human osteoarthritis cartilage by global gene expression analysis. Osteoarthritis Cartilage 26, 1531–1538. 10.1016/j.joca.2018.07.012.

100. Rintala, T.J., Ghosh, A., and Fortino, V. (2022). Network approaches for modeling the effect of drugs and diseases. Brief Bioinform 23. 10.1093/bib/bbac229.

101. Langfelder, P., and Horvath, S. (2008). WGCNA: an R package for weighted correlation network analysis. BMC Bioinformatics 9, 559. 10.1186/1471-2105-9-559.

102. Liu, X., Liu, R., Croker, B.A., Lawlor, K.E., Smyth, G.K., and Wicks, I.P. (2015). Distinctive pro-inflammatory gene signatures induced in articular chondrocytes by oncostatin M and IL-6 are regulated by Suppressor of Cytokine Signaling-3. Osteoarthritis Cartilage 23, 1743–1754. 10.1016/j.joca.2015.05.011.

103. Mobasheri, A., Saarakkala, S., Finnilä, M., Karsdal, M.A., Bay-Jensen, A.C., and van Spil, W.E. (2019). Recent advances in understanding the phenotypes of osteoarthritis. F1000Res 8. 10.12688/f1000research.20575.1.

104. Cao, Y., Tang, S.a., and Ding, C. (2021). Inflammatory phenotype of osteoarthritis and its potential therapies. Rheumatology & Autoimmunity 1, 92–100. https://doi.org/10.1002/rai2.12018.

105. Yuan, C., Pan, Z., Zhao, K., Li, J., Sheng, Z., Yao, X., Liu, H., Zhang, X., Yang, Y., Yu, D., et al. (2020). Classification of four distinct osteoarthritis subtypes with a knee joint tissue transcriptome atlas. Bone Res 8, 38. 10.1038/s41413-020-00109-x.

106. Subramanian, A., Narayan, R., Corsello, S.M., Peck, D.D., Natoli, T.E., Lu, X., Gould, J., Davis, J.F., Tubelli, A.A., Asiedu, J.K., et al. (2017). A Next Generation Connectivity Map: L1000 Platform and the First 1,000,000 Profiles. Cell 171, 1437–1452.e1417. 10.1016/j.cell.2017.10.049.

107. Lv, Z., Xu, X., Sun, Z., Yang, Y.X., Guo, H., Li, J., Sun, K., Wu, R., Xu, J., Jiang, Q., et al. (2021). TRPV1 alleviates osteoarthritis by inhibiting M1 macrophage polarization via Ca(2+)/CaMKII/Nrf2 signaling pathway. Cell Death Dis 12, 504. 10.1038/s41419-021-03792-8.

108. Nakagome, K., Imamura, M., Okada, H., Kawahata, K., Inoue, T., Hashimoto, K., Harada, H., Higashi, T., Takagi, R., Nakano, K., et al. (2011). Dopamine D1-like receptor antagonist attenuates Th17-mediated immune response and ovalbumin antigen-induced neutrophilic airway inflammation. J Immunol 186, 5975–5982. 10.4049/jimmunol.1001274.

109. Alleboina, S., Wong, T., Singh, M.V., and Dokun, A.O. (2020). Inhibition of protein kinase C beta phosphorylation activates nuclear factor-kappa B and improves postischemic recovery in type 1 diabetes. Exp Biol Med (Maywood) 245, 785–796. 10.1177/1535370220920832.

110. Ağaç, D., Estrada, L.D., Maples, R., Hooper, L.V., and Farrar, J.D. (2018). The β2-adrenergic receptor controls inflammation by driving rapid IL-10 secretion. Brain Behav Immun 74, 176–185. 10.1016/j.bbi.2018.09.004.

111. Bonnans, C., Chou, J., and Werb, Z. (2014). Remodelling the extracellular matrix in development and disease. Nat Rev Mol Cell Biol 15, 786–801. 10.1038/nrm3904.

112. Iijima, H., Gilmer, G., Wang, K., Bean, A.C., He, Y., Lin, H., Tang, W.Y., Lamont, D., Tai, C., Ito, A., et al. (2023). Age-related matrix stiffening epigenetically regulates α-Klotho expression and compromises chondrocyte integrity. Nat Commun 14, 18. 10.1038/s41467-022-35359-2.

113. Alford, J.W., and Cole, B.J. (2005). Cartilage restoration, part 1: basic science, historical perspective, patient evaluation, and treatment options. Am J Sports Med 33, 295–306. 10.1177/0363546504273510.

114. van der Kraan, P.M., Buma, P., van Kuppevelt, T., and van den Berg, W.B. (2002). Interaction of chondrocytes, extracellular matrix and growth factors: relevance for articular cartilage tissue engineering. Osteoarthritis Cartilage 10, 631–637. 10.1053/joca.2002.0806.

115. Yamamoto, R., Chung, R., Vazquez, J.M., Sheng, H., Steinberg, P.L., Ioannidis, N.M., and Sudmant, P.H. (2022). Tissue-specific impacts of aging and genetics on gene expression patterns in humans. Nat Commun 13, 5803. 10.1038/s41467-022-33509-0.

116. Wang, X., Jiang, Q., Song, Y., He, Z., Zhang, H., Song, M., Zhang, X., Dai, Y., Karalay, O., Dieterich, C., et al. (2022). Ageing induces tissue-specific transcriptomic changes in Caenorhabditis elegans. Embo j 41, e109633. 10.15252/embj.2021109633.

117. Mahmoudian, A., Lohmander, L.S., Mobasheri, A., Englund, M., and Luyten, F.P. (2021). Early-stage symptomatic osteoarthritis of the knee - time for action. Nat Rev Rheumatol 17, 621–632. 10.1038/s41584-021-00673-4.

118. Sanchez-Lopez, E., Coras, R., Torres, A., Lane, N.E., and Guma, M. (2022). Synovial inflammation in osteoarthritis progression. Nat Rev Rheumatol 18, 258–275. 10.1038/s41584-022-00749-9.

119. Wojdasiewicz, P., Poniatowski, Ł.A., and Szukiewicz, D. (2014). The role of inflammatory and anti-inflammatory cytokines in the pathogenesis of osteoarthritis. Mediators of inflammation 2014.

120. van Beuningen, H.M., Arntz, O.J., and van den Berg, W.B. (1991). In vivo effects of interleukin-1 on articular cartilage. Prolongation of proteoglycan metabolic disturbances in old mice. Arthritis Rheum 34, 606–615. 10.1002/art.1780340513.

121. Latourte, A., Kloppenburg, M., and Richette, P. (2020). Emerging pharmaceutical therapies for osteoarthritis. Nat Rev Rheumatol 16, 673–688. 10.1038/s41584-020-00518-6.

122. Banerjee, S., Biehl, A., Gadina, M., Hasni, S., and Schwartz, D.M. (2017). JAK-STAT Signaling as a Target for Inflammatory and Autoimmune Diseases: Current and Future Prospects. Drugs 77, 521–546. 10.1007/s40265-017-0701-9.

123. Li, W.Q., Dehnade, F., and Zafarullah, M. (2001). Oncostatin M-induced matrix metalloproteinase and tissue inhibitor of metalloproteinase-3 genes expression in chondrocytes requires Janus kinase/STAT signaling pathway. J Immunol 166, 3491–3498. 10.4049/jimmunol.166.5.3491.

124. Ji, T., Chen, M., Sun, W., Zhang, X., Cai, H., Wang, Y., and Xu, H. (2022). JAK-STAT signaling mediates the senescence of cartilage-derived stem/progenitor cells. J Mol Histol. 10.1007/s10735-022-10086-6.

125. Denton, C.P., Del Galdo, F., Khanna, D., Vonk, M.C., Chung, L., Johnson, S.R., Varga, J., Furst, D.E., Temple, J., Zecchin, C., et al. (2022). Biological and clinical insights from a randomized phase 2 study of an anti-oncostatin M monoclonal antibody in systemic sclerosis. Rheumatology (Oxford) 62, 234–242. 10.1093/rheumatology/keac300.

126. Wright, H.L., Moots, R.J., and Edwards, S.W. (2014). The multifactorial role of neutrophils in rheumatoid arthritis. Nat Rev Rheumatol 10, 593–601. 10.1038/nrrheum.2014.80.

127. Mimpen, J.Y., Baldwin, M.J., Cribbs, A.P., Philpott, M., Carr, A.J., Dakin, S.G., and Snelling, S.J.B. (2021). Interleukin-17A Causes Osteoarthritis-Like Transcriptional Changes in Human Osteoarthritis-Derived Chondrocytes and Synovial Fibroblasts In Vitro. Front Immunol 12, 676173. 10.3389/fimmu.2021.676173.

128. Kriegová, E., Manukyan, G., Mikulková, Z., Gabcova, G., Kudelka, M., Gajdos, P., and Gallo, J. (2018). Gender-related differences observed among immune cells in synovial fluid in knee osteoarthritis. Osteoarthritis and cartilage 26, 1247–1256.

129. Hsueh, M.F., Zhang, X., Wellman, S.S., Bolognesi, M.P., and Kraus, V.B. (2021). Synergistic roles of macrophages and neutrophils in osteoarthritis progression. Arthritis & Rheumatology 73, 89–99.

130. Wang, G., Jing, W., Bi, Y., Li, Y., Ma, L., Yang, H., and Zhang, Y. (2021). Neutrophil Elastase Induces Chondrocyte Apoptosis and Facilitates the Occurrence of Osteoarthritis via Caspase Signaling Pathway. Front Pharmacol 12, 666162. 10.3389/fphar.2021.666162.

131. Moran, E.M., Mullan, R., McCormick, J., Connolly, M., Sullivan, O., Fitzgerald, O., Bresnihan, B., Veale, D.J., and Fearon, U. (2009). Human rheumatoid arthritis tissue production of IL-17A drives matrix and cartilage degradation: synergy with tumour necrosis factor-alpha, Oncostatin M and response to biologic therapies. Arthritis Res Ther 11, R113. 10.1186/ar2772.

132. Franceschi, C., Garagnani, P., Parini, P., Giuliani, C., and Santoro, A. (2018). Inflammaging: a new immune-metabolic viewpoint for age-related diseases. Nat Rev Endocrinol 14, 576–590. 10.1038/s41574-018-0059-4.

133. Supek, F., Bošnjak, M., Škunca, N., and Šmuc, T. (2011). REVIGO summarizes and visualizes long lists of gene ontology terms. PloS one 6, e21800.

134. Greene, C.S., Krishnan, A., Wong, A.K., Ricciotti, E., Zelaya, R.A., Himmelstein, D.S., Zhang, R., Hartmann, B.M., Zaslavsky, E., and Sealfon, S.C. (2015). Understanding multicellular function and disease with human tissue-specific networks. Nature genetics 47, 569–576.

135. Robinson, M.D., McCarthy, D.J., and Smyth, G.K. (2010). edgeR: a Bioconductor package for differential expression analysis of digital gene expression data. bioinformatics 26, 139–140.

136. Ritchie, M.E., Phipson, B., Wu, D., Hu, Y., Law, C.W., Shi, W., and Smyth, G.K. (2015). limma powers differential expression analyses for RNA-sequencing and microarray studies. Nucleic acids research 43, e47–e47.

137. Moher, D., Liberati, A., Tetzlaff, J., and Altman, D.G. (2009). Preferred reporting items for systematic reviews and meta-analyses: the PRISMA statement. Ann Intern Med 151, 264–269, w264. 10.7326/0003-4819-151-4-200908180-00135.

138. Shamseer, L., Moher, D., Clarke, M., Ghersi, D., Liberati, A., Petticrew, M., Shekelle, P., Stewart, L.A., and Group, P.-P. (2015). Preferred reporting items for systematic review and meta-analysis protocols (PRISMA-P) 2015: elaboration and explanation. BMJ 349, g7647. 10.1136/bmj.g7647.

139. Stroup, D.F., Berlin, J.A., Morton, S.C., Olkin, I., Williamson, G.D., Rennie, D., Moher, D., Becker, B.J., Sipe, T.A., and Thacker, S.B. (2000). Meta-analysis of observational studies in epidemiology: a proposal for reporting. Meta-analysis Of Observational Studies in Epidemiology (MOOSE) group. Jama 283, 2008–2012. 10.1001/jama.283.15.2008.

140. Higgins, J.P., and Green, S. (2011). Cochrane handbook for systematic reviews of interventions (John Wiley & Sons).

141. Vesterinen, H.M., Sena, E.S., Egan, K.J., Hirst, T.C., Churolov, L., Currie, G.L., Antonic, A., Howells, D.W., and Macleod, M.R. (2014). Meta-analysis of data from animal studies: a practical guide. Journal of neuroscience methods 221, 92–102. 10.1016/j.jneumeth.2013.09.010.

142. Methley, A.M., Campbell, S., Chew-Graham, C., McNally, R., and Cheraghi-Sohi, S. (2014). PICO, PICOS and SPIDER: a comparison study of specificity and sensitivity in three search tools for qualitative systematic reviews. BMC Health Serv Res 14, 579. 10.1186/s12913-014-0579-0.

143. Jackson, S.J., Andrews, N., Ball, D., Bellantuono, I., Gray, J., Hachoumi, L., Holmes, A., Latcham, J., Petrie, A., Potter, P., et al. (2017). Does age matter? The impact of rodent age on study outcomes. Laboratory animals 51, 160–169. 10.1177/0023677216653984.

144. Little, C.B., and Hunter, D.J. (2013). Post-traumatic osteoarthritis: from mouse models to clinical trials. Nat Rev Rheumatol 9, 485–497. 10.1038/nrrheum.2013.72.

145. Rongen, J.J., Hannink, G., van Tienen, T.G., van Luijk, J., and Hooijmans, C.R. (2015). The protective effect of meniscus allograft transplantation on articular cartilage: a systematic review of animal studies. Osteoarthritis and cartilage 23, 1242–1253. 10.1016/j.joca.2015.04.025.

146. Bricca, A., Juhl, C.B., Steultjens, M., Wirth, W., and Roos, E.M. (2019). Impact of exercise on articular cartilage in people at risk of, or with established, knee osteoarthritis: a systematic review of randomised controlled trials. Br J Sports Med 53, 940–947. 10.1136/bjsports-2017-098661.

147. Pritzker, K.P., Gay, S., Jimenez, S.A., Ostergaard, K., Pelletier, J.P., Revell, P.A., Salter, D., and van den Berg, W.B. (2006). Osteoarthritis cartilage histopathology: grading and staging. Osteoarthritis and cartilage 14, 13–29. 10.1016/j.joca.2005.07.014.

148. Mankin, H.J., Dorfman, H., Lippiello, L., and Zarins, A. (1971). Biochemical and metabolic abnormalities in articular cartilage from osteo-arthritic human hips. II. Correlation of morphology with biochemical and metabolic data. The Journal of bone and joint surgery. American volume 53, 523–537.

149. Moussavi-Harami, S.F., Pedersen, D.R., Martin, J.A., Hillis, S.L., and Brown, T.D. (2009). Automated objective scoring of histologically apparent cartilage degeneration using a custom image analysis program. J Orthop Res 27, 522–528. 10.1002/jor.20779.

150. Jørgensen, A.E.M., Kjær, M., and Heinemeier, K.M. (2017). The Effect of Aging and Mechanical Loading on the Metabolism of Articular Cartilage. J Rheumatol 44, 410–417. 10.3899/jrheum.160226.

151. López-López, J.A., Page, M.J., Lipsey, M.W., and Higgins, J.P.T. (2018). Dealing with effect size multiplicity in systematic reviews and meta-analyses. Research synthesis methods. 10.1002/jrsm.1310.

152. Rohatgi, A. (2015). WebPlotDigitizer (Version 3.9). http://arohatgi.info/WebPlotDigitizer.

153. Drevon, D., Fursa, S.R., and Malcolm, A.L. (2017). Intercoder Reliability and Validity of WebPlotDigitizer in Extracting Graphed Data. Behav Modif 41, 323–339. 10.1177/0145445516673998.

154. Koo, T.K., and Li, M.Y. (2016). A Guideline of Selecting and Reporting Intraclass Correlation Coefficients for Reliability Research. J Chiropr Med 15, 155–163. 10.1016/j.jcm.2016.02.012.

155. Bolia, I.K., Mertz, K., Faye, E., Sheppard, J., Telang, S., Bogdanov, J., Hasan, L.K., Haratian, A., Evseenko, D., Weber, A.E., and Petrigliano, F.A. (2022). Cross-Communication Between Knee Osteoarthritis and Fibrosis: Molecular Pathways and Key Molecules. Open Access J Sports Med 13, 1–15. 10.2147/oajsm.S321139.

156. A single-cell transcriptomic atlas characterizes ageing tissues in the mouse. (2020). Nature 583, 590-595. 10.1038/s41586-020-2496-1.

157. Deeks, J.J., and Higgins, J.P. (2010). Statistical algorithms in review manager 5. Statistical Methods Group of The Cochrane Collaboration, 1-11.

158. Higgins, J.P., Thompson, S.G., Deeks, J.J., and Altman, D.G. (2003). Measuring inconsistency in meta-analyses. BMJ 327, 557–560. 10.1136/bmj.327.7414.557.

159. van Middelkoop, M., Arden, N.K., Atchia, I., Birrell, F., Chao, J., Rezende, M.U., Lambert, R.G., Ravaud, P., Bijlsma, J.W., Doherty, M., et al. (2016). The OA Trial Bank: meta-analysis of individual patient data from knee and hip osteoarthritis trials show that patients with severe pain exhibit greater benefit from intra-articular glucocorticoids. Osteoarthritis Cartilage 24, 1143–1152. 10.1016/j.joca.2016.01.983.

160. Wold, S., Esbensen, K., and Geladi, P. (1987). Principal component analysis. Chemometrics and intelligent laboratory systems 2, 37–52.

161. Chen, Y., Lun, A.T., and Smyth, G.K. (2016). From reads to genes to pathways: differential expression analysis of RNA-Seq experiments using Rsubread and the edgeR quasi-likelihood pipeline. F1000Res 5, 1438. 10.12688/f1000research.8987.2.

